# Panx1 Ablation Aggravates Oxidative Stress and Cell Death by Altering AMPK/mTOR signaling Pathways and the Composition of Synapses in the Zebrafish

**DOI:** 10.1101/2024.04.23.590821

**Authors:** Georg S.O. Zoidl, Nickie Safarian, Christiane Zoidl, Steven Connor, Georg R. Zoidl

## Abstract

Pannexin-1 channels have garnered attention for their implications in neurodevelopment, potentially having a dual role in mediating a delicate balance between cell death and survival. However, a comprehensive understanding of the underlying molecular and cellular mechanisms and Panx1’s potential protective functions throughout neurodevelopment remains to be determined. Zebrafish larvae with loss of Pannexin-1a function were subjected to an acute exposure to 1-methyl-4-phenyl-1,2,3,6-tetrahydropyridine (MPTP). Early-life changes in larvae induced by uncoupling of oxidative phosphorylation were investigated by a computational Gene Set Expression Analysis of RNA-seq data and experimental testing of light-stimulated locomotor behavior, cell death, and bioelectrical properties of local field potentials in the ascending visual pathway. A KEGG pathway analysis underscored Panx1a’s regulatory influence on neurodevelopment. Targeting Panx1a caused a deregulation of oxidative phosphorylation, glycolysis, reactive oxygen production, hypoxia, unfolded protein response pathways, and reduced extracellular ATP. Further, Panx1a ablation enhanced the transcriptional activation of 5’ AMP-activated protein kinase (AMPK) kinase, a cellular energy sensor activated by falling energy status, largely to activate glucose and fatty acid uptake and oxidation when cellular energy is low. The activation of the AMPK pathway in Panx1a knock-out larvae correlated with the stimulation of the mammalian target of rapamycin (mTORC1) pathway that controls cellular metabolism, catabolism, immune responses, autophagy, survival, proliferation, and migration, to maintain cellular homeostasis. The differential expression of mTORC1 pathway genes associated with autophagy, and apoptosis signaling pathways. The resultant cell death was pronounced in the pallium and tectum regions. The loss of cells interrelated with a trans-synaptic a loss of synaptic neurotransmitter receptor and ion channel/transporter expression. Local field potential recordings in the optic tectum and pallium demonstrated that Panx1a’s involvement in modulating local neuronal networks was altered. Collectively, the results shed light on the impacts of acute MPTP treatment on locomotor behavior, transcriptomic shifts, metabolic disturbances, and the pivotal role of Panx1a in cell death. These insights enhance our comprehension of the intricate molecular and cellular mechanisms underpinning neurodevelopment, with implications for potential therapeutic strategies targeting Panx1 channels in autism and Alzheimer’s disease.

**Graphical Abstract:** 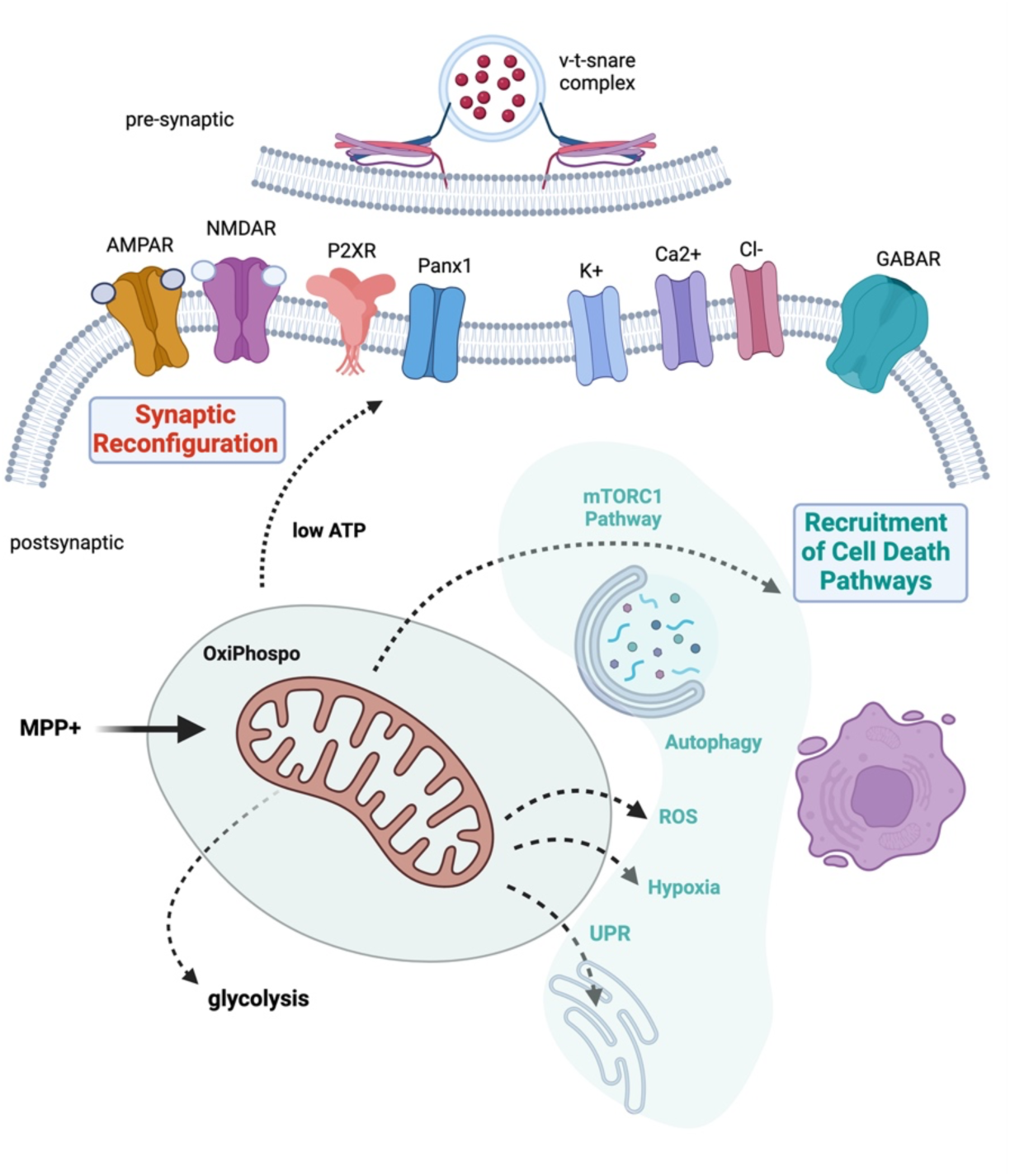

**Highlights:**

- A genetic model was combined with the neurotoxin MPTP to explore the roles of zebrafish Pannexin-1 channels in neurodevelopment under oxidative stress conditions.
- A potential beneficial impact of targeting Panx1a is superseded by synaptic plasticity loss, dysfunctional mitochondrial metabolism, and cell death pathway activation.
- Loss of Panx1a amplifies AMPK/mTORC1 pathway activation of cell death pathways.
- A role of Panx1a as a regulator of energy and synaptic homeostasis was identified.

## Introduction

Pannexin 1 (Panx1) forms channels with significant roles in neurodevelopment, synapse function, and cellular signaling and communication between neurons and glial cells. The multifaceted roles of this channel in brain development encompasses structural aspects of neuronal and synaptic function such as the formation and stabilization of dendritic spines, which are essential for synaptic connectivity and plasticity. The absence of Panx1 leads to higher dendritic spine densities and more intricate neuronal networks, indicating that Panx1 regulates the dynamics of dendritic and spiny protrusions, thus influencing the overall architecture of neuronal networks.

In addition to structural roles, Panx1 channels are key modulators of synaptic transmission and plasticity. For example, Panx1 knockout (Panx1^-/-^) mice exhibit enhanced excitability and more complex dendritic branching. These changes are linked to increased actin polymerization and imbalances in Rho GTPase activities, suggesting that Panx1 interacts with the cytoskeleton to regulate synapse function. Panx1 also regulates long-term potentiation (LTP) and long-term depression (LTD) at hippocampal synapses, processes critical for learning and memory. Panx1^-/-^ mice display deficiencies in long-term spatial reference memory, implicating Panx1 in memory consolidation.

Panx1’s role in neuron-glia interactions and ATP release is crucial for neuro-developmental processes. It is vital for neural stem cell survival, neuronal maturation, and synaptic plasticity. Furthermore, Panx1’s role in ATP release and purinergic signaling is crucial for neurodevelopmental processes, including NPC proliferation and migration, which are foundational for establishing synchronized neuronal networks and preventing pathological conditions such as seizures. A recently published Panx1 interactome reveals associations with proteins involved in synaptic function and neurodevelopmental and neurodegenerative diseases, indicating a broader role in synaptic homeostasis and network functions.

Overall, Panx1 has emerged as a critical player in brain development, influencing synaptic formation, stability, plasticity, neuron-glia communication, and neuronal network formation. The functions extend to maintaining synaptic stability, contributing to memory and learning, and participating in signaling pathways essential for brain function. While Panx1’s involvement in neuronal development and system circuitry formation is documented, the underlying molecular and cellular mechanisms of how these channels influence embryogenesis and early postnatal development await a more detailed characterization.

Zebrafish larvae are an established model for studying cell signaling and communication during embryogenesis and early postnatal development. The extra-corporal embryogenesis ends with the hatching of transparent larvae at three days post fertilization (3dpf). At six days post fertilization (6dpf), during post-embryonic development, extensive cellular communication drives growth, differentiation, and responses to environmental cues. The rapid development of behaviors signifies the maturation of functional neuronal networks.

We were specifically interested in the role of dopaminergic neurons during this period due to their extensive roles in developmental processes regulating the formation and maturation of neural circuits, influencing behaviors, and cognitive functions. Furthermore, imbalances of dopaminergic signaling potentially lead to conditions such as autism, attention-deficit/hyperactivity disorder (ADHD), and schizophrenia, or present as a risk factor for neurodegenerative disorders (NDDs) like Parkinson’s disease (PD).

We have previously identified in 6dpf zebrafish larvae a neurodevelopmental deficit when ablation of Panx1a, the ortholog of mammalian Panx1, caused visuomotor deficiency linked to changes in dopaminergic signaling. To get insight into the roles of Panx1 in dopaminergic signaling, we targeted dopaminergic neurons by inducing rapid neurotoxicity in six-day-old Tubingen Longfin (TL) larvae with 1-methyl-4-phenyl-1,2,3,6-tetrahydropyridine (MPTP). MPTP crosses the blood-brain barrier and is converted to the toxic MPP+ by the mitochondrial monoamine oxidase (Mao-B). MPP+ is taken up by the dopamine transporter (DAT) expressed by dopaminergic neurons, inactivating the mitochondrial respiratory complex I. By disrupting dopaminergic signaling in a Panx1 knockout zebrafish line (Panx1a^-/-^) we tested the hypothesis that Panx1a acts as a regulator linking mitochondrial health with synapse function. The computational analysis of RNA-seq data showed that a four-hour treatment in TL (Panx1a^+/+^) and Panx1a^-/-^ larvae was sufficient to initiate transcriptome-wide expression changes indicating metabolic dysfunction, impaired proteostasis, differential expression of genes involved in neurodevelopmental and neurodegenerative disorders, and cell death. MPTP also caused a dose-dependent reduction in swimming behavior. By comparing MPTP effects on Panx1a^-/-^ and Panx1a^+/+^ strains, RNA-seq, and Gene Set Expression Analysis (GSEA), we found that Panx1a ablation altered the expression of hallmark genes for autism, Alzheimer’s, and Parkinson’s disease, reducing the expression of zebrafish orthologs of mammalian Shank, alpha-Synuclein, and Tau proteins. In contrast, Panx1a ablation reduced extracellular ATP, and enhanced differential expression of cell signaling pathways regulating oxidative phosphorylation, cell death, lipid synthesis, ROS signaling, and intracellular quality controls. Ablation of Panx1a also altered synapse molecular composition, impairing synaptic function, demonstrated *in vivo* by recording changes to local field potentials. Alterations of in the optic tectum (OT) and dorso-medial (DM) pallium corroborated neuronal network dysfunctions.

In summary, this study’s findings suggest an intricate relationship between Panx1a, synapses, and mitochondrial health underscoring the importance of maintaining mitochondrial integrity and balanced dopamine signaling for optimal neurodevelopment. The animal model introduced here can provide insights into potential therapeutic targets for mitigating the impact of these disorders on neurodevelopmental health.

## Materials and Methods

### Zebrafish lines

Zebrafish (*Danio Rerio*) of strain Tubingen long fin (TL) were maintained in groups with mixed sex in a recirculation system (Aquaneering Inc., San Diego, CA) at 28⁰C on a 14hr light/10hr dark cycle. All animal work was performed at York University’s zebrafish vivarium and in an S2 biosafety laboratory following the Canadian Council for Animal Care guidelines after approval of the study protocol by the York University Animal Care Committee (GZ#2019-7-R2). The *Panx1a* knockout line (*panx1a^-/-^*) was generated and characterized in-house ^1–3^.

### Zebrafish Locomotion Assays

The Zebrabox behavior system and the ZebraLab analytical suite (ViewPoint Life Technology, Lyon, France, http://www.viewpoint.fr) were used for automated extraction of behavioral outputs and video tracking. Tracking videos were recorded at 30 frames per second (fps) under infrared (for Light-OFF recording) or visible light (Light-ON) illumination using a Point Grey Research Dragonfly2 DR2-HIBW camera (Teledyne-FLIR, Burlington, ON, Canada). Inside the Zebrabox a lightbox provided visible and infrared light from below for recordings up to 40% light intensity (visible range, 0-1200 lux). 6dpf larvae were observed in clear 48-well plates maintained at 28⁰C throughout the experiment. All experiments were performed between 12:00 to 2 pm, as larvae (6 dpf) activity was previously reported to reach a stable level by early afternoon ^4^. Locomotor activity scores (total distance ± SEM.; n = 16 per group) were determined from larvae’s movements in the time window from 2100 to 2300 seconds. Panx1a^+/+^ and Panx1a^-/-^ larvae were exposed to MPTP (Sigma-Aldrich) for four hours prior to recording. The Zebrabox provided a Light-OFF stimulus for 20 minutes at 0% intensity (0 lux), followed by incremental 10% light intensity increases every 10 minutes up to 40% (1200 lux). A principal component analysis (PCA) was performed as described ^1,5^. The statistical analysis of locomotion data used a Welch’s t-test.

### *In vivo* electrophysiology

Published procedures were used to prepare anaesthetized 6 days post fertilization (6dpf) zebrafish larvae for *in vivo* electrophysiology ^1,6^. Zebrafish larvae at 6dpf were anesthetized using 0.3mM Pancuronium bromide (Sigma-Aldrich, Oakville, ON, Canada) for 2-3 min until the touch response stopped. Anesthetized larvae were immobilized in freshly prepared 2% low melting temperature agarose. A Leica S9E dissecting microscope (Leica Microsystems, Richmond Hill, ON, Canada) was used to orient the dorsal aspect of the larvae to the gel surface. Embedded larvae were placed on the upright stage of an Olympus BX51 fluorescence microscope (Olympus, Richmond Hill, ON, Canada). Larvae were submerged in 1ml of egg water (E3; pH 7.2-7.4) that was applied topically to the agar. Under direct visual guidance two glass microelectrodes (1.2mM OD, approximately 1µM tip diameter, 2-7MΩ), backloaded with 2M NaCl, were placed into the right optic tectum (OT) and the dorsomedial (DM) region of the pallium. Local field potentials were recorded simultaneously from both microelectrodes using a Multiclamp 700B amplifier (Axon Instruments, San Jose, CA, USA). Voltage recordings were low-pass filtered at 1kHz (-3 dB; eight-pole Bessel), high-pass filtered at 0.1 Hz, digitized at 10 kHz using a Digidata 1550A A/D interface, and stored on a PC running pClamp11 software (all Axon Instruments). The basal activity was recorded for 10 minutes under Light-ON conditions (1000 lux), during which images of the electrode placement were taken for reference. For each fish, brain activity was normalized to its baseline activity to account for the biological variability of individual brains. In the treatment group larvae were exposed to MPTP (50µM) four hours before recording. A temperature of 28⁰C was maintained during experiments.

### Power spectral density and coherence detection analysis

A Power spectral density (PSD) estimation was performed by the Welch’s method for baseline and MPTP recordings. A moving window for fast Fourier transform (FFT) computation was used and all windows were averaged. The frequency bands theta (3.5-7.5 Hz), beta (12-30 Hz) and gamma (35-45 Hz) were extracted for analysis using NeuroExplorer Version 4 (Nex Technologies, Colorado Springs, CO, 80906, USA). Changes in PSD between baseline and treatment were analyzed for significance using a Welch’s t-test. A Coherence analysis was performed applying the Hann method for baseline and MPTP recordings using NeuroExplorer. Changes in the coherence between baseline and treatment were tested for significance using a Welch’s t-test.

### Extracellular ATP Assay

For quantification of ATP and protein ≈ 50 6dpf larvae were collected at experimental endpoints. Larvae were homogenized in 500µl ice-cold of Phosphate-Buffered Saline (PROBS) containing 100µM ARL-67156 (Sigma-Aldrich) and Halt Protease inhibitor (1:100) (Thermo-Fisher Scientific, Waltham, Massachusetts, USA) for 1min at 30Hz using the Tissuelyser^LT^ (Qiagen, Hilden, Germany). Homogenates were transferred to chilled Eppendorf tubes (^-^20⁰C) and centrifuged at 12,000 rpm for 2min. Supernatants were collected and snap-frozen in liquid nitrogen before storage at ^-^80⁰C. Extracellular ATP measurements used a 96-well format (Greiner Bio-One, Monroe, North Carolina, USA) and the Molecular PROBes^®^ ATP determination Kit as described by the manufacturer (Life Technologies, Burlington, ON, Canada). ATP was quantified in replicates of 6 using the Synergy H4 Hybrid Multi-well Plate Reader (Biotek, Winooski, Vermont, USA). The luminescent assay parameters were: plate temperature set to 28⁰C; a low-intensity shake of the plate for 3s before reading; a 5s integration time per well; gain setting at 150. In every test, the ATP concentration in experimental samples was calculated from ATP standard curves (0-1µM ATP) included in the same 96-well plate. Data were exported from the Gen5 Data Analysis Software (Biotek) and analyzed in Excel and GraphPad Prism 10. ATP was represented as a normalized concentration per mg of protein. A NanoDrop Spectrophotometer (Thermo Fisher Scientific) was used to measure protein content in 2µL of supernatant from homogenized samples. The final ATP content was expressed after normalizing the protein content in each sample of pooled larvae ^7^. The data sets were compared using Welch’s t-test.

### Cell Death Assay

Up to seven larvae (6dpf) were placed into a 2ml Eppendorf tube in egg water. MPTP treatment (20µM) was applied for four hours. During the last 60mins, acridine orange (2.5µl/ml; Sigma-Aldrich) was added to the Eppendorf tube. Following incubation, the larvae freely swam in 10ml fresh egg water for 10 minutes to remove excess AO. Then, the larvae were immobilized using the muscle relaxant pancuronium bromide and mounted head down in 2% agar noble in dishes. Cell death was imaged using a Zeiss LSM700 (Objective 10x NA0.8; Z-stack ≈ 100µm). All parameters were controlled by ZEN software (ZEISS, North York, ON, Canada) and consistently applied. Raw 3D volumes were processed for AO-positive cell detection using Bitplane-Imaris software (Oxford Instruments, Abingdon, UK). Statistics used unpaired t-test and GraphPad Prism 10.

### NGS RNA-sequencing

The transcriptomes of all zebrafish lines used in this research were analyzed by RNA-seq (NGS-Facility, The Center for Applied Genomics, SickKids, Toronto, ON, Canada). The reported data was derived from the sequencing of three independent pools of ≈30 age-matched larvae (6dpf). The RNA-seq data are deposited at the NCBI -Gene Expression Omnibus (GEO) database repository (ID pending). Total RNAs were extracted from 30 6dpf larvae using RNeasy Plus Mini Kit (Qiagen). The RNA quality was determined using a Bioanalyzer 2100 DNA High Sensitivity chip (Agilent Technologies, Mississauga, ON, Canada). The RNA library preparation was performed following the NEB NEBNext Ultra II Directional RNA Library Preparation protocol (New England Biolabs Inc., Ipswich, MA, USA). RNA libraries were loaded on a Bioanalyzer 2100 DNA High Sensitivity chip to check for size, quantified by qPCR using the Kapa Library Quantification Illumina/ABI Prism Kit protocol (KAPA Biosystems, Wilmington, MA, USA). Pooled libraries were paired-end sequenced on a High Throughput Run Mode flow cell with the V4 sequencing chemistry on an Illumina HiSeq 2500 platform (Illumina, Inc., San Diego, CA) following Illumina’s recommended protocol to generate paired-end reads of 126-bases in length.

### Differential Gene Expression and Functional Enrichment Analyses

The post-sequencing processing to final read counts, normalization, and differential gene expression analysis used multiple software packages, including a two-condition differential expression analysis using the edgeR R-package, v.3.8.6 ^8,9^ and DESeq2 R package version 1.40.2 ^10^. Genotypes and treatments were incorporated into the statistical model and multiple hypothesis testing was performed. The default filter for DESeq2 used a threshold of p<0.05 and the Benjamini-Hochberg procedure to determine the false discovery rate (FDR), and the adjusted P-value (padj). Differential gene expression was defined as log2FoldChange >2 or <-2, respectively.

A Gene Set Enrichment Analysis (GSEA) tested whether a defined set of genes shows statistically significant, concordant differences between genotypes and treatment groups. This method calculates sample-wise gene set enrichment scores and estimates variation of pathway activity over a sample population in an unsupervised manner ^10^. The Molecular Signatures Database (MSigDB) v.5.2 (http://software.broadinstitute.org/gsea/msigdb/index.jsp), a renowned repository of curated gene sets encompassing diverse biological pathways was used to source gene sets for broad Hallmarks. Before GSEA RNA-seq data were filtered at padj < 0.01 to increase the stringency of the analysis. Other tools used to process RNA-seq data were HCOP (https://www.genenames.org/tools/hcop/) and db2db at bioDBnet (https://biodbnet.abcc.ncifcrf.gov/db/db2db.php) to convert curated human or mouse to zebrafish genes ^11,12^. Curated gene lists were generate with information from the Zebrafish Information Network (ZFIN) (https://zfin.org/) and the KEGG (Kyoto Encyclopedia of Genes and Genomes, https://www.genome.jp/kegg/) databases, and using unique qualifiers from the Gene Ontology (GO) project (https://geneontology.org/).

### Pharmacology

All chemicals were purchased from Sigma-Aldrich (Mississauga, Canada): (MPTP; 1-50µM; cat#: M0896), Pancuronium bromide (Panc; 300µM; cat#P1918), Acridine-Orange (AO; 15µM; cat#: A6014). All reagents were water-soluble. The concentrations referred to in results are final.

### Statistics and data reproducibility

Statistical analyses were performed in NeuroExplorer VS4, or in GraphPad Prism VS10. Results are represented as the mean ± standard error of the mean (SEM). For molecular analysis (RNA-seq, ATP detection), a minimum of n ≥ 3 independent experimental replicates were generated. For behavioral testing G*power analysis determined the number of larvae ^1,5^. The normality and homogeneity of the data variance were determined by the “Shapiro-Wilk Test” and “Levene’s Test.” Experimental groups were compared using unpaired t-tests or Welch’s t-tests. A *P* value <0.05 was considered statistically significant. For all experiments, sample sizes, statistical tests, and when appropriate *P* values are indicated in the figure legends.

## Results

### Acute MPTP treatment the locomotion of zebrafish larvae

Acute MPTP treatment for four hours was evaluated to establish a practical alternative of targeting mitochondrial metabolism in dopaminergic neurons during a critical developmental stage. The six-days post fertilization (6dpf) timepoint was chosen based on the expression profile of Panx1 genes in rodents and fish; the expression reaches a transient peak before birth in rodents ^13^, and when zebrafish transition to free-swimming juveniles.

To assess the impact of MPTP treatment, we utilized a modified visual-motor response (VMR) assay, measuring zebrafish larvae responsiveness to gradual Light-ON changes up to 1200 lux after a period of Light-Off (**Fig 1a**). A principal component analysis (PCA) of 11 swimming behavioral parameters, captured over 72% (PC1) and 24% (PC2) of the data variance, respectively. The total swim distance (PC1, *totaldist*) was used to quantify differences between MPTP-treated and untreated genotypes (**Fig. 1b**).

**Figure 1:**
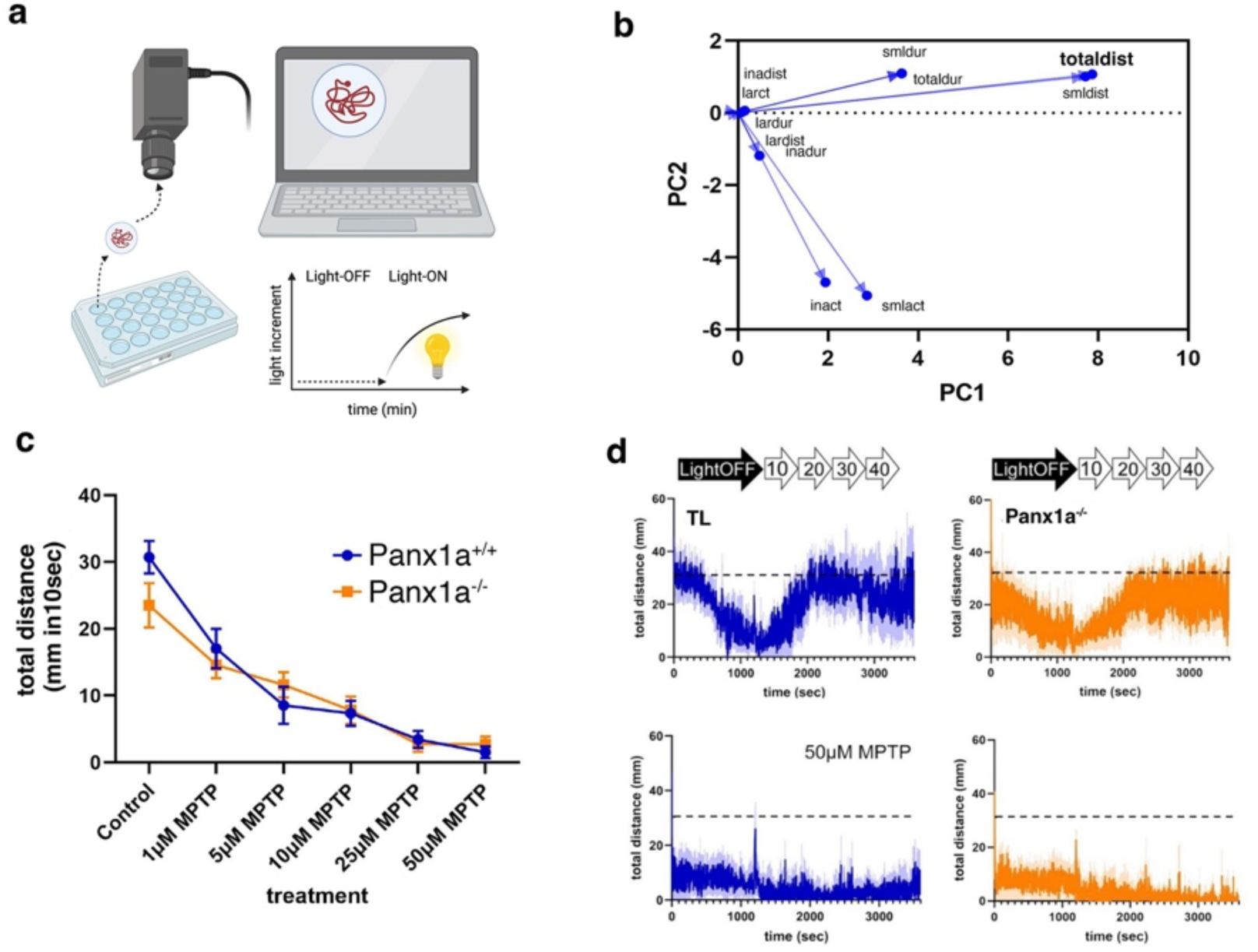
Acute MPTP treatment alters zebrafish locomotion. a) Experimental setup for video recording and light stimulation of 6dpf larvae during a modified visual-motor response (VMR) assay. b) Principal Component Analysis of 11 behavioral parameters. c) Concentration-dependent impact of acute MPTP treatment on swimming behavior without (control) or after 4hrs exposure. d) Cumulative traces representing the Mean (dark blue/orange) and SEM (light blue/orange) for n=16 larvae/genotype and MPTP treatment condition. The horizontal dashed line indicates the average total swimming distance of untreated Panx1a^+/+^ larvae. Statistics: Student t-test; Values represent Mean ± SEM for total distance of swimming in mm in a 10-second time-bin. The image in a) was created with BioRender.

In the VMR assay, both Panx1a^+/+^ and Panx1a^-/-^ larvae responded to MPTP treatment (1µM to 50µM) in Light-ON conditions, with a decline in swimming distance as the MPTP dose increased (**Fig. 1c**). Acute MPTP treatment consistently decreased swimming activity in both Panx1a^+/+MPTP^ and Panx1a^-/-MPTP^ larvae compared to untreated controls, even at the lowest concentration tested. The average total distance covered in a 10-second time bin was 25.6 ± 0.8 mm (Panx1a^+/+^) and 21.1 ± 0.4 mm (Panx1a^-/-^), with no statistically significant differences between genotypes in the presence of MPTP. **Fig. 1d** shows that the initial Light-OFF period reduced swimming distance to a baseline, followed by the resumption of swimming activity in untreated larvae as light increased from 10% (400 lux) to 40% (1200 lux) Light-ON over the next 40 minutes. This result underscores the impact of Panx1a ablation and acute MPTP treatment on the visuomotor response in 6dpf zebrafish larvae.

### Panx1a^+/+MPTP^ and Panx1a^-/-MPTP^ larvae show different transcriptional responses associated with altered mitochondrial metabolism, cell homeostasis, and cell death

To gain insight into the molecular basis of the observed behavioral changes experimental RNA-seq analysis and a computational approach compared MPTP-treated Panx1a knockout (Panx1a^-/-MPTP^) and wild-type (Panx1a^+/+MPTP^) larvae with naïve siblings, unraveling key regulatory pathways underlying MPTP neurotoxicity in the model. Gene expression profiles of Panx1a^-/-^ and Panx1a^+/+^ larvae were different before MPTP exposure; RNA-seq identified a total of 31251 individual RNA transcripts, with n=724 genes upregulated and n=1030 genes downregulated in the knockout group compared to wild-type controls **(Fig.2a)**. Following the exposure to MPTP (4 hours), both genotypes exhibited substantial alterations in gene expression patterns, highlighting the dynamic nature of the cellular response to MPTP. Notably, the magnitude of gene expression dysregulation was markedly amplified in Panx1a^-/-MPTP^ larvae post-treatment compared to Panx1a^+/+MPTP^ counterparts (downregulated: Panx1a^+/+MPTP^ n=3764 genes, Panx1a^-/-MPTP^ n=4828 genes; upregulated: Panx1a-^+/+MPTP^ n=3500 genes, Panx1a^-/-MPTP^ n=4749 genes; false discovery rate: FDR < 0.05) (**Fig. 2b,c**).

**Figure 2:**
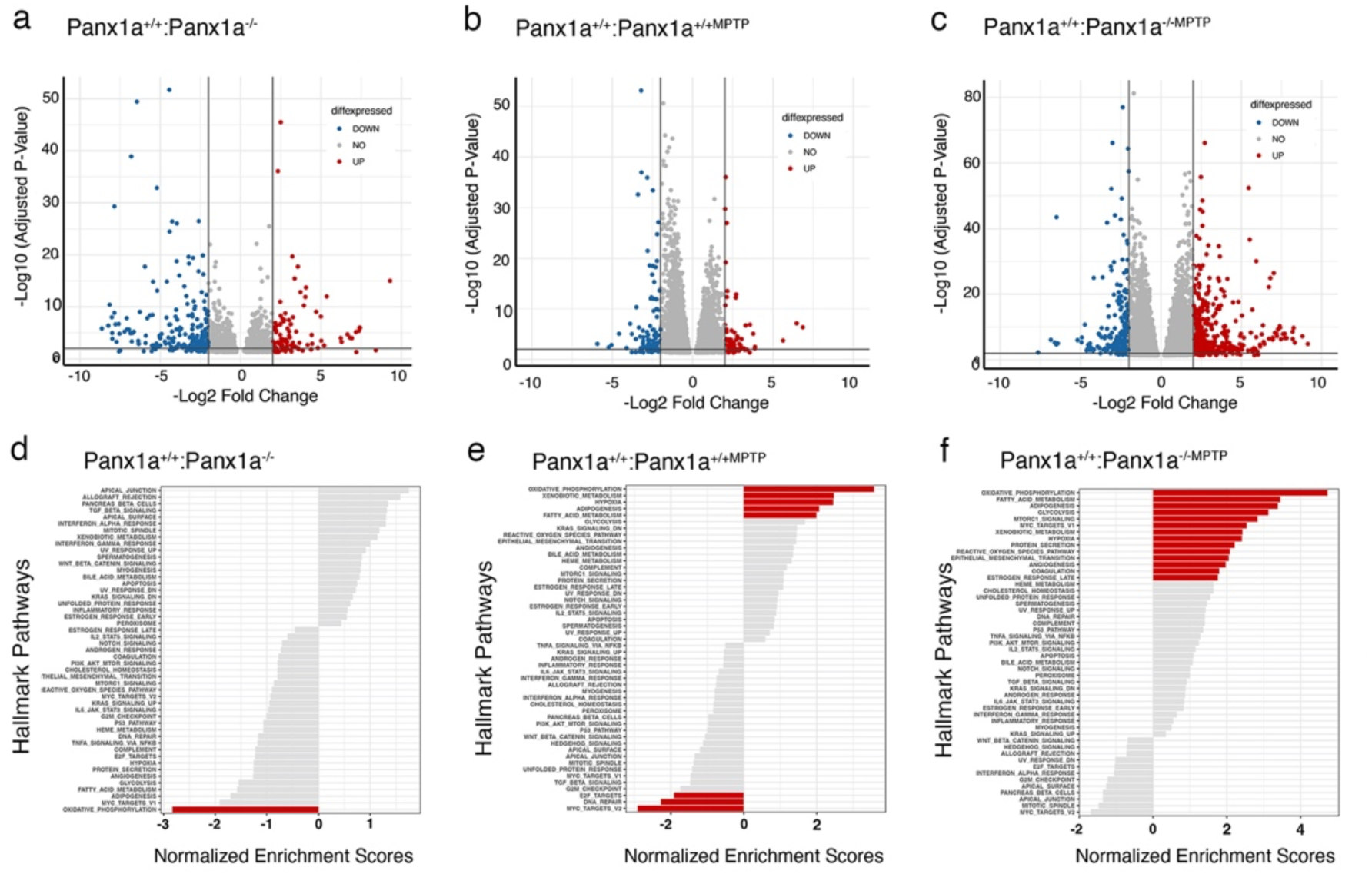
MPTP treatment caused greater disruption of gene expression in Panx1a^-/-^ larvae. a-c) illustrates the log2 fold change of differentially expressed genes pre-and post-MPTP exposure, with changes exceeding a two-fold threshold highlighted in red (upregulated) or blue (downregulated). d-f) Gene set enrichment analysis (GSEA) revealing normalized enrichment scores (NES) for the top 50 ranked Hallmark pathways. Enriched pathways are denoted in red (threshold: padj <0.05), indicating noteworthy alterations in biological processes associated with MPTP treatment in knock-out larvae compared to wild-type counterparts.

To interpret the biological pathways perturbed by MPTP treatment, we conducted a gene set enrichment analysis (GSEA) ^14^. The visualization of enrichment within the top 50 Hallmarks pathways provided first insights into intricate regulatory networks modulating neuronal responses to MPTP exposure in the context of Panx1a ablation. Our analysis revealed significant enrichment of pathways associated with metabolism and cell death, including oxidative phosphorylation (Oxiphospho), glycolysis, reactive oxygen species (ROS) regulation, mammalian target of rapamycin complex 1 (mTORC1) signaling, fatty acid metabolism, and hypoxia. Considering the mitochondrial dysfunction in MPTP-induced neurotoxicity, the next step of our investigation prioritized the analysis of how Panx1a ablation affected pathways linked to mitochondrial homeostasis.

### The MPTP-induced metabolic crisis altered pathways regulating hypoxia, reactive oxygen species, and fatty acid homeostasis

A comprehensive computational analysis, leveraging GSEA, explored the impact of MPTP treatment on Panx1a^+/+^ and Panx1a^-/-^ zebrafish larvae’s mitochondrial health. The GSEA result demonstrated an enrichment of the oxidative phosphorylation pathway in both Panx1a^+/+MPTP^ (n=100) and Panx1a^-/-MPTP^ larvae (n=155) with a log2 fold filter set to >0.5 and <-0.5 and a padj of <0.01 (**Fig. 3a**). Only n=38 genes were different in untreated controls showing that MPTP treatment impacted this pathway significantly.

**Figure 3:**
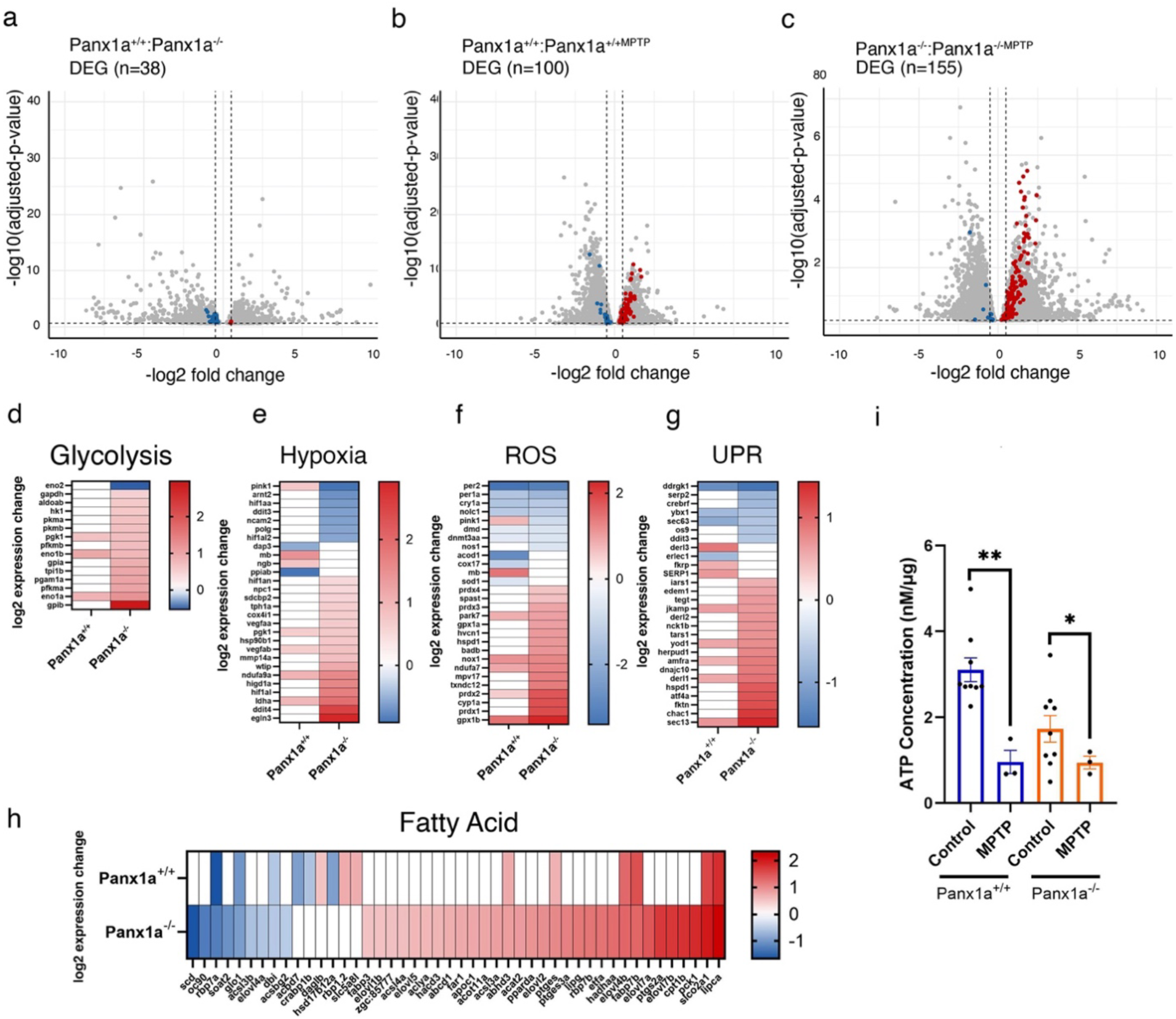
Acute treatment with MPTP caused a metabolic crisis and reduced ATP release in Panx1a^+/+^ larvae. a-c) Differentially expressed genes in the oxidative phosphorylation pathway. Dots in blue and red represent significantly regulated genes in the oxidative phosphorylation pathway. Grey dots represent all DEGs. d-h) Heatmaps illustrate the differential expression of genes identified in the biological processes: GO:0006096 – *Glycolysis*, GO:0001666 -*Response to Hypoxia*, GO:0000302 - *Response to reactive oxygen species* (ROS), GO:0030968 - *Unfolded Protein Response (UPR)*, and GO:0006631 - *Fatty acid metabolic process*. i) ATP concentrations were measured before and after acute MPTP treatment. MPTP treatment caused a significant downregulation in Panx1a^+/+MPTP^ larvae (P-value < 0.05). No significant differences in ATP concentrations were observed in Panx1a^-/-MPTP^ larvae. The ATP concentrations were adjusted for protein concentration to evaluate extracellular ATP levels across multiple samples. Each sample represented a pool of 50 6dpf larvae. Statistical approach: Welch’s t-test. The number of experimental replicates: Panx1a^+/+^ and Panx1a^-/-^ untreated controls n=9; MPTP treatment conditions n=3. Mean ± SEM. Statistical significance was indicated as *: P-value * <0.05, ns = not significant.

The notable differential gene expression modulation and directionality of changes in Panx1a^-/-MPTP^ larvae was also found in biological processes downstream of oxidative phosphorylation, such as *GO:0006096* – *Glycolysis*, *GO:0001666 - Response to Hypoxia*, *GO:0000302 - Response to reactive oxygen species (ROS), GO:0030968 - Unfolded Protein Response (UPR), and GO:0006631 - Fatty acid metabolic process* (**Fig. 3d-h**; **Suppl. Tables 2- 6)**.

Notable Panx1a^-/-MPTP^ specific regulations of glycolysis genes were the dysregulation of two enolase genes, *eno1a/b* and *eno2*, key enzymes for cellular bioenergetics, without which glycolysis cannot produce ATP, which reduces hypoxia tolerance (**Fig. 3d**, **Suppl. Table 2)**. Likewise, the differential expression of pyruvate kinases *pkma/b*, phosphofructokinases *pfkma/b*, and phosphoglycerate mutase A (*pgam1a*) demonstrate the switch to an oxygen-independent, and relatively inefficient way of ATP generation when Complex I is blocked by MPTP treatment.

Furthermore, MPTP treatment caused the differential expression of hypoxia or ROS related pathways in Panx1a^-/-MPTP^ larvae that can promote neuronal cell death (*hif1al, ldha, ddit4, egln3*) **(Fig. 3 e,f, Suppl. Tables 3,4)**. A notable change to expression in the unfolded protein response (UPR) was a gene encoding for a protein, *sec13*, that functions as an activator of the amino acid-sensing branch of the mTORC1 signaling pathway (**Fig. 3g**; **Suppl. Table 5)**. Other regulated genes like *fktn, chac1, atf4a* are part of a signaling complex linked to synaptic function. More specifically, the mammalian homolog of *fktn* regulates synaptic function via inhibition of the ATF4 transcription factors complex with DISC1 (Disrupted-in-schizophrenia 1), or function like *chac1*as a pro-apoptotic component of the unfolded protein response pathway by mediating the pro-apoptotic effects of the ATF4 in the ATF4-ATF3-DDIT3/CHOP cascade.

The notable upregulation of *Fatty acid metabolic process* (GO:0006631) illustrated how Panx1a ablation affected energy homeostasis (**Fig. 3h**; **Suppl. Table 6)**. For example, the upregulated fatty acid elongation enzymes represented by *elovl1b, elovl4b, elovl5,* or *elovl7a/b* are important for the mitochondrial fatty acid β-oxidation, the major pathway for the degradation of fatty acids that is essential for maintaining energy homeostasis. The transcriptional regulation of five apolipoproteins *apoea/b, apoa1a, apobb.1*, and *apoc1* signified pleiotropic effects on lipoprotein metabolism. The upregulation of *apoea/b* was notable since human ortholog(s) are implicated in multiple diseases, including Alzheimer’s disease, artery disease, cerebrovascular disease, eye disease, and familial hyperlipidemia. We reasoned that an accumulation of fatty acids in mitochondria above physiological levels could have harmed mitochondrial function in Panx1a^-/-MPTP^ larvae causing the observed changes to hypoxia and reactive oxygen species-related processes.

We hypothesized that the altered expression of genes in the oxidative phosphorylation pathway affected ATP production, with the absence of the ATP-release channel Panx1a amplifying the effect of Complex I blockage by MPP+. To validate this hypothesis, we quantified extracellular ATP concentrations both before and after MPTP treatment in both genotypes. The impact of MPTP treatment on extracellular ATP levels was pronounced in Panx1a^+/+MPTP^ larvae, demonstrating a significant reduction (Panx1a^+/+^: 3.12 ± 0.28 nM/µg; Panx1a^+/+MPTP^: 0.96 ± 0.27 nM/µg; P-value=0.0011), indicating a limitation in ATP production (**Fig. 3i**). Conversely, MPTP treatment had a minor effect on Panx1a^-/-MPTP^ larvae, revealing a decrease of low ATP levels (Panx1a^-/-^: 1.73 ± 0.31 nM/µg; Panx1a^-/-MPTP^: 0.95 ± 0.15 nM/µg; P-value=0.044).

The upregulation of mitochondrial Complex I and Complex III genes, NADH:ubiquinone oxidoreductase subunit A11 (*ndufa11*), or ubiquinol-cytochrome c oxidoreductases (*uqcr10, uqcrfs1*), illustrate changes to mitochondrial energy metabolism and homeostasis which are amplified in Panx1a^-/-MPTP^ larvae. The augmented upregulation of pathways leading to hypoxia, ROS-production, or the unfolded protein response further substantiate that Panx1a is required to maintain metabolic and energetic homeostasis *in vivo*.

### Loss of Panx1a and activation of oxidative phosphorylation alters the AMPK-mTOR pathway

The enrichment of the mTORC1 hallmark, identified through GSEA (see Fig. 2), implied mechanistically a switch towards anabolic metabolism. The differential gene expression of mTORC1 pathway genes in untreated controls (n=13), Panx1^+/+MPTP^ (n= 74), and Panx1a^-/-MPTP^ larvae (n=121) exhibited an enrichment trend that showed that Panx1a^-/-MPTP^ were affected most by the oxidative phosphorylation (OXIPHOS) dysfunction (**Fig. 4a-c**). The upregulation of the AMP-activated protein kinase (*prkaa1*/AMPK), a master regulator of cellular energy homeostasis, together with genes in the AMPK pathway in Panx1a^-/-MPTP^ larvae further suggested that a molecular response to stresses that deplete cellular ATP supplies such as Complex I blockage was found (**Fig. 4a-c**).

**Figure 4:**
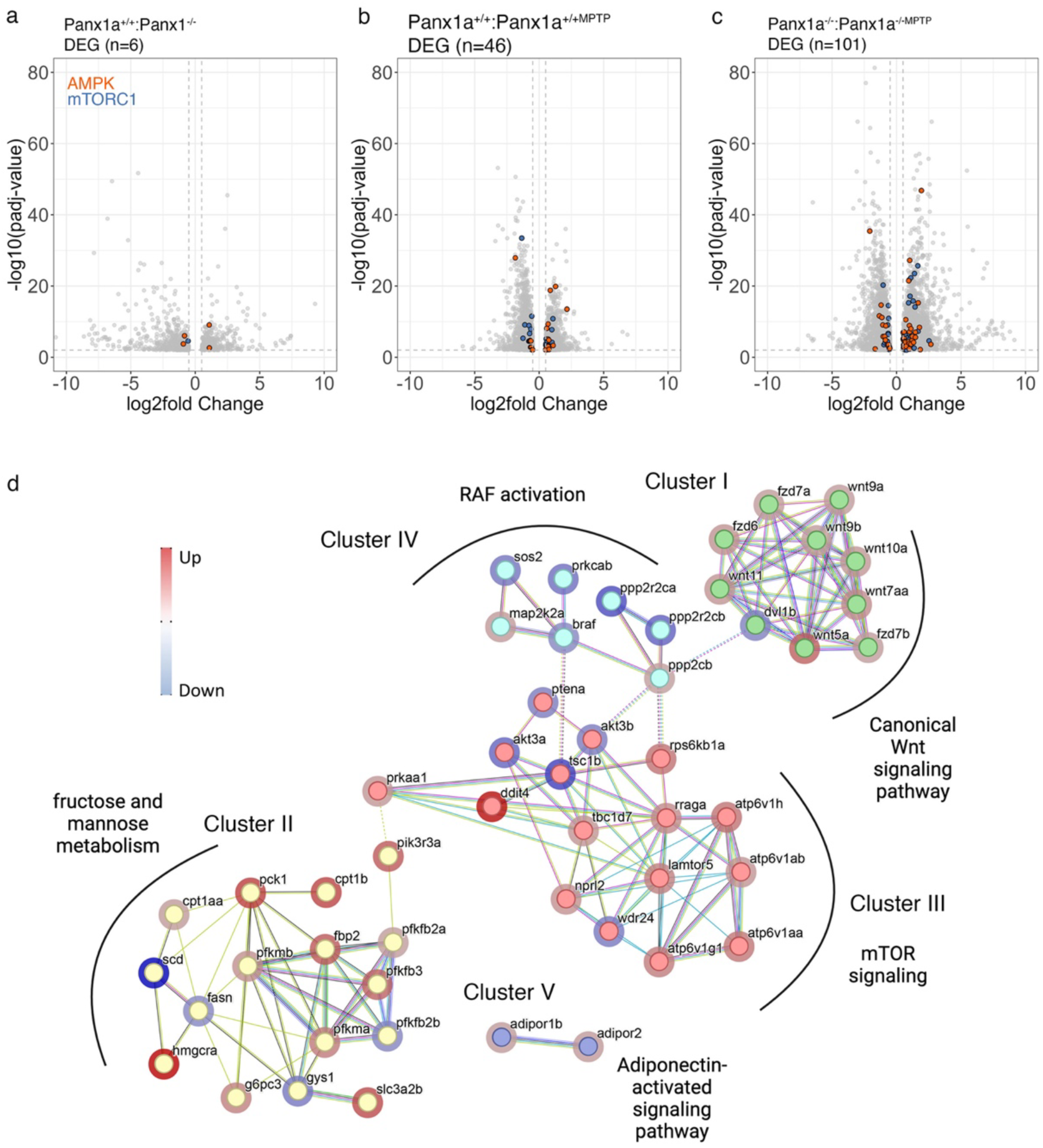
Loss of Panx1a and activation of oxidative phosphorylation alters the AMPK-mTOR pathway. a-c) Vulcano plots represent the differential expression of AMPK-(orange) and mTORC1 (blue) pathway genes. d) STRING analysis of 49 genes. K-means clustering identified five clustered groups.

To identify specific molecular pathways in response to stress when cellular ATP is depleted in Panx1a^-/-MPTP^ larvae we used STRING analysis to infer context (**Fig. 4d**). The input were 54 AMPK-mTOR pathway genes, ranked by their log2fold expression. K-means clustering identified five clusters specific for Panx1a^-/-MPTP^ larvae. Notable groups of genes upregulated in the mTOR pathway included nine Wnt/frizzled genes in Cluster I (*wnt5a, wnt7aa, wnt9a/b, wnt10a, wnt11, fdz6, fdz7a/b*). Activation of the canonical Wnt-signaling pathway in neurons could enhance both hexose utilization through glycolysis and regulate dendrite formation and complexity. Indeed, Cluster II encompasses 16 genes representing fructose and mannose metabolism. Cluster II linked to Cluster III through upregulated *prkaa1a/AMPK* and *pik3r3a.* The 1-phosphatidylinositol-3-kinase regulator activity acts upstream of the insulin receptor signaling pathway and the phosphatidylinositol phosphate biosynthetic process. Within Cluster III the upregulation of four subunits of vacuolar ATPases (*V-ATPase; atp6v1aa atp6v1ab atp6v1b2, atp6v1g1 atp6v1h atp6v1c1b atp6v1ba atp6v1f*) implicated a disruption of presynaptic cellular homeostasis in neurons. Furthermore, late endosomal/lysosomal adaptor and MAPK and mTOR activator genes (*lamtor5*) and Ras-related GTP-binding protein A (*rraga*) encode proteins forming the Lamtor/Ragulator complex that regulates multiprotein signaling units on late endosomes/lysosomes with crucial roles in the cellular response to amino acid availability through regulation of the mTORC1 signaling cascade.

The downregulation of RAF activation (Cluster IV) indicated the complexity of molecular pathway changes affected by Panx1a ablation in cell stress. A reduction of RAF activation as an entry point to the mitogen-activated protein kinase (MAPK)/extracellular signal-regulated kinase (ERK-1/2) signaling pathway, suggested effects on fundamental cellular functions including proliferation, differentiation, and survival.

The adiponectin signaling pathway, and the Adiponectin activated signaling pathway (Cluster 5) involved two receptors that are required for normal glucose and fat homeostasis. In mice AdipoR1 and AdipoR2 are present in the synaptosome, with AdipoR2 displaying increased presynaptic vs. postsynaptic localization, whereas AdipoR1 was enriched in both the presynaptic and postsynaptic fractions. Loss of function mutations in mouse KO models support transsynaptic roles causing cognitive deficits.

### Loss of Panx1a and activation of oxidative phosphorylation promotes cell death pathways

The enrichment of the mTORC1 pathway observed in the GSEA analysis suggested responses to metabolic and other stresses. A follow-up of the differential expression regulation of 192 mTOR pathway genes identified 121 regulated genes in Panx1a^-/-MPTP^ larvae. Fewer genes were differentially regulated in Panx1a^+/+^ (n=14) and Panx1a^+/+MPTP^ ^larvae^ (n=74) (**Fig. 5a**).

**Figure 5:**
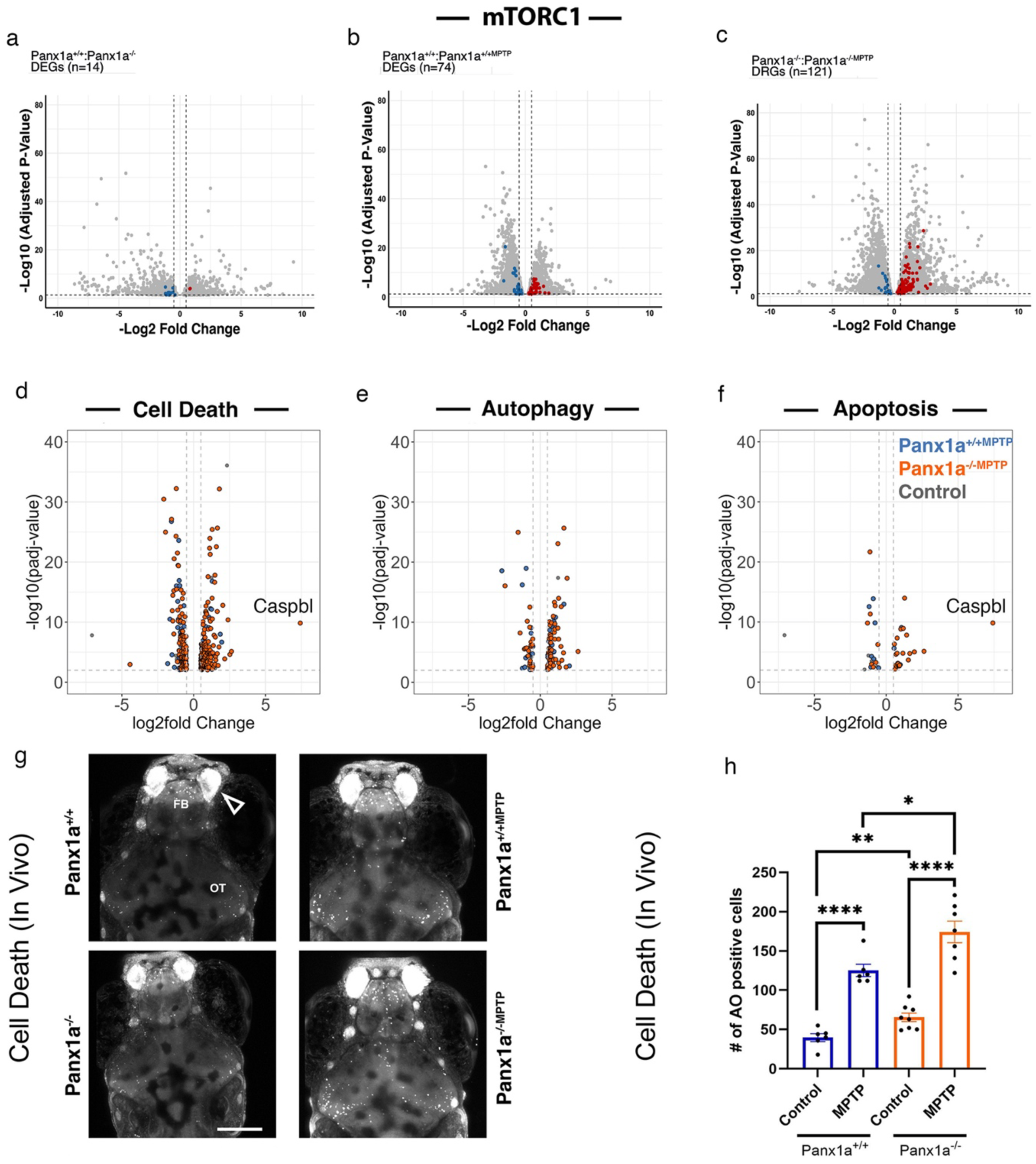
Loss of Panx1a promotes cell death in the acute MPTP model. a-c) Differentially regulated genes in the mTORC1 pathway. Blue and red dots represent regulated genes in the pathway. Grey dots represent all DEGs. d-f) Significantly regulated genes (padj ≤ 0.01) were compared against zfin lists of cell death, apoptosis, and autophagy. Regulated genes were plotted. g) Experimental outline of the *in-vivo* cell death assay using the acridine orange dye. Representative projection views of Z-stacks representing a depth ≈ 100µm collected from untreated and MPTP-treated Panx1a^+/+^ and Panx1a^-/-^ larvae. h) Quantification of acridine-orange positive cells. Statistical approach: Significance was calculated using an unpaired t-test. Number of larvae, n=7 for all treatments. Mean ± SEM. Statistical significance was indicated as *: P-values * <0.05, ** <0.01, *** <0.005, **** <0.001.

The upregulation of eight subunits of vacuolar ATPases (*V-ATPase; atp6v1aa atp6v1ab atp6v1b2, atp6v1g1 atp6v1h atp6v1c1b atp6v1ba atp6v1f*) and late endosomal/lysosomal adaptor and MAPK and mTOR activator genes (LAMTOR/Ragulator) encoding scaffold proteins forming complexes that regulate multiprotein signaling units on late endosomes/lysosomes were involved in acidification (*lamtor2, lamtor3, lamtor5*). We propose that the upregulation of these components could reflect an attempt by the cell to adapt to stress by enhancing lysosomal function and autophagy.

These changes suggested that systems in MPTP-related larvae in the absence of Panx1a were overwhelmed or dysfunctional due to an inability to maintain cellular homeostasis, leading to cell death. The functional enrichment analysis showed differentially expressed genes associated with cell death (GO:0008219), autophagy (GO:0006914), and the execution phase of apoptosis (GO:0097194). In Panx1a^-/-MPTP^ larvae, 223 cell death-related genes were identified, while Panx1a^+/+MPTP^ larvae exhibited regulation in 107 genes (**Fig 5b**, **Suppl Table 7)**. The analysis of autophagy genes mirrored the overall pattern of cell death, with a substantial six to eleven-fold increase in differentially expressed genes when comparing untreated with MPTP-treated larvae (Panx1a^+/+MPTP^, n=38; Panx1a^-/-MPTP^, n=70) (**Fig. 5c**). The unique molecular response in the absence of Panx1a was evident, with Panx1a^-/-MPTP^ larvae showing a notable increase in differentially upregulated (n=35) and downregulated genes (n=16) **(Suppl Table 8)**. Among the upregulated genes, transmembrane 41b (*tmem41b*), tumor protein p53-inducible nuclear protein 1 (*tp53inp1*), and the stimulator of interferon (*sting1*), pointed towards the activation of autophagy and cellular stress response. These genes play essential roles in autophagosome biogenesis, pro-apoptotic responses, and inflammation mediation, respectively, contributing to cellular adaptations and responses under MPTP-induced conditions.

Activation of the execution phase of apoptosis was observed in Panx1a^-/-MPTP^ larvae, as demonstrated by the upregulation of seven members of the caspase gene family, including initiators of apoptosis *casp9* and *casp20*, and executioner phase caspases *casp3a, casp6a, casp6b1*, and *casp7*. The log2 fold ratio = 7.15 upregulation of the inflammatory caspase *caspbl* (alternative name casp19b) was particularly notable as a hallmark for the execution phase of apoptosis, downstream of the assembly of the inflammasome complex, leading to the cleavage and activation of the proinflammatory cytokine *il1b* (**Fig. 5f**, **Suppl Table 9)**.

An *in-vivo* cell death stain, acridine orange (AO), was chosen as a direct way of visualizing cell death (**Fig. 5g**). Notably, AO-stained nuclei were particularly prominent in brain regions known for dopaminergic neuron expression or the optic tectum, which receives inputs from hypothalamic dopaminergic neurons. In the absence of treatment, Panx1a^+/+^ larvae exhibited the lowest number of dead cells (Panx1a^+/+^ = 39.67, Panx1a^-/-^ = 66). The number of AO-positive cells in Panx1a^-/-^ larvae was significantly higher compared to Panx1a^+/+^ (P-value = 0.0056) (**Fig. 5h**). MPTP treatment increased cell death in both genotypes (P-value = 0.0130), with an average of n=174 AO-positive cells in Panx1a^-/-MPTP^ and n=125 in Panx1a^+/+MPTP^ larvae.

In summary, the results underscore a multifaceted role for Panx1a in modulating gene expression related to cell death, apoptosis, and autophagy. Imaging of cell death *in vivo* implicates regions with loss of dopaminergic neurons.

### Trans-synaptic changes after Panx1a ablation and MPTP treatment

A DESeq2 analysis of the RNA sequencing analysis, employing a log2 gene filter of >0.5 and <-0.5 with a padj < 0.01, identified 681 differentially expressed genes in Panx1^-/-MPTP^ and Panx1^+/+MPTP^ larvae. The analysis of the Gene Ontology search terms *GO:Biological Process* and *GO:Cellular Compartment* indicated significant upregulation of biosynthetic pathways, suggestive of increased metabolic activities in 6dpf larvae with Panx1a ablation (**Fig. 6a,b**). Conversely, downregulated genes were enriched for biological processes related to neurotransmission at chemical synapses, synaptic signaling, and neurodevelopment, affecting both pre-and postsynaptic compartments. Specifically, the query term *GO:0045202 – Synapse* identified 53 downregulated genes and no upregulated genes, while the terms *GO:0098793 – Presynapse* and *GO:0098794 – Postsynapse* identified 35 and 36 differentially expressed genes, respectively (**Fig. 6c,d**; **Suppl Table 10)**.

**Figure 6:**
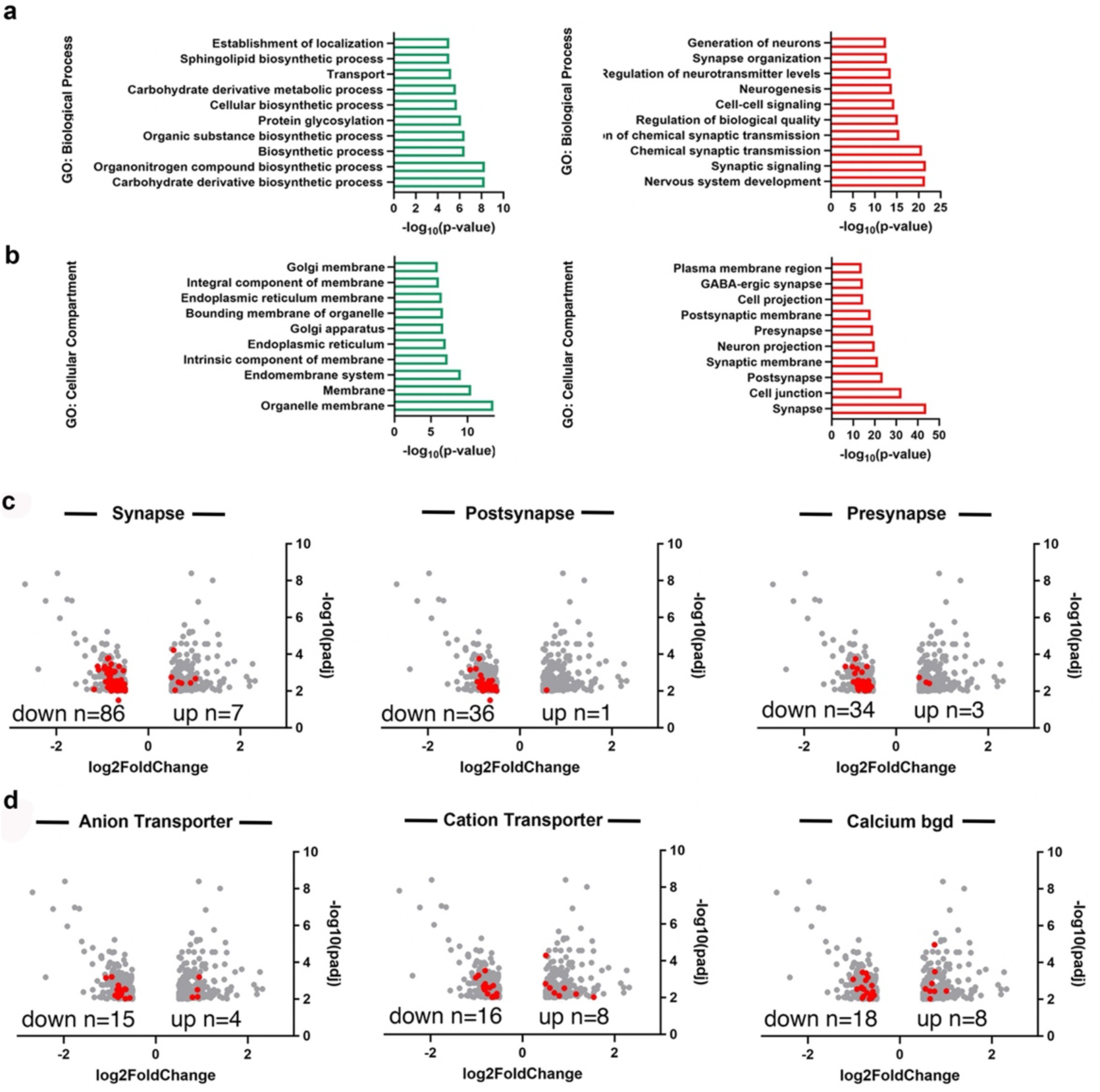
Trans-synaptic changes after Panx1a ablation and MPTP treatment. a,b) Summary of the ten most enriched Gene Ontology Biological Processes and Cellular compartments found in Panx1a^+/+MPTP^ and Panx1a^-/-MPTP^ treated larvae. The P-values of each category represent the false discovery rate presented as inverse log10. c), d) Volcano plots represent the differentially expressed genes in Panx1a^+/+MPTP^ and Panx1a^-/-MPTP^ treated larvae (red dots). Gray dots represent the 671 differentially expressed genes found in both genotypes. The cutoff filter was: log2 >0.5 or <-0,5 and padj <0.01.

STRING analysis of the downregulated genes demonstrated a significant impact of Panx1a ablation on GABAergic and glutamatergic transmission. Notably, ligand-gated and G protein-coupled GABA receptor subtypes *(gabra6b, gabrb1, gabrb3, gabrb4, gabrz, gabrg2, gabrr2a)* (**Cluster I)**, ionotropic NMDA (*grin2aa, grin2ab, grin2bb*), AMPA (*gria4a*), and kainite receptors (*grik2*) were downregulated alongside other regulators of synaptic plasticity (**Cluster II Suppl. Fig. 1**). For example, the mouse ortholog of *shisa9a* is known as a regulator of short-term neuronal synaptic plasticity in the dentate gyrus associated with AMPA receptor function. Likewise, the discs large MAGUK scaffold protein genes *dlg2* (PSD93) and *dlg4* (PSD95) were downregulated.

**Cluster III** encompassed genes encoding pre-synaptic proteins with essential roles in neurotransmitter release via the formation of the soluble N-ethylmaleimide-sensitive fusion protein attachment protein receptors (SNARE) complex (synaptosomal-associated protein: *snap25a, snap25b*), vesicle exocytosis or calcium dynamics in response to depolarization (synaptotagmin: *syt1a, syt5b*; syntaxin: *stx1b, stx12l, stxbp1a*; complexin; *cplx2, cplx2l, cplx3b, cplx4a*). The presynaptic downregulation of glutamate decarboxylase gene 2 (*gad2*) expression is notable; the Gad2 protein catalyzes the presynaptic production of GABA, which suggests a negative feedback loop as downregulation of GABA production corresponds with reduced GABA receptor expression.

Furthermore, deregulation of inorganic ion transporters (n=45) and calcium binding proteins (n=27) was observed in Panx1a^-/-MPTP^ larvae (**Fig. 6d**). Synaptically localized anionic ion transporters (n=15) with reduced expression in the nervous system **(Suppl. Fig 1, Clusters I, II)** included AMPA and NMDA receptors, or chloride permeable GABA receptors. The four up-regulated genes included a Na-Cl cotransporter-like protein (*slc12a10.2*), an anion:anion antiporter with bicarbonate transmembrane transporter activity (*slc4a1b*), or a gene encoding a voltage-gated chloride channel (*clcn2c*) that maintains chloride ion homeostasis. Deregulation of cation transporters (n=25) included transporters and channels permeable for potassium (*kcna1b, kcnip1b, kcnip3b, kcn1a.1, kcnk3b*), calcium (*cacna1fb, cacna1ha*), sodium (*scn3b*), or co-transporters for sodium and chloride (*slc12a10.2*), symporters for potassium and chloride (*slc12a5b*), a potassium-dependent sodium/calcium exchanger (*slc24a5*), or a GABA:sodium:chloride symporter (*slc6a1a*). The loss of Panx1a also affected the differential expression of calcium-binding proteins (n=26). Notable examples include members of the protocadherin gene family (*pcdh1a, pcdh8, pcdh17, pcdh19*) regulating neurotransmission and synaptic plasticity.

Overall, the computational analysis suggested that Panx1a ablation, in conjunction with MPTP treatment, perturbed ion homeostasis and neurotransmitter release at synapses, affecting neurotransmission and synaptic plasticity across pre-and postsynaptic compartments.

### Loss-of-Panx1a function alters the power of theta, beta-and gamma-waves in the optic tectum

The impact of MPTP-induced cell death and alterations in energy metabolism on brain activity was assessed by Local Field Potential (LFP) recordings. We determined how the absence of Panx1a influenced local neuronal circuits following acute MPTP treatment by strategically positioned electrodes in the optic tectum (OT) and dorsomedial forebrain (DM), regions previously identified as having cell loss after exposure to 10µM MPTP (**Fig. 7a**). Simultaneous recordings of brain waves were performed on awake, immobilized 6dpf larvae. We categorized frequency spectra into specific bands (Theta: 3.5-7.5Hz, Beta: 12-30Hz, Gamma: 35-45Hz), employing a common approach used in mammalian electrophysiological data analysis that is applicable to zebrafish. Raw traces (**Fig. 7b**) underwent lowpass filtering (<300Hz) to isolate Theta, Beta, and Gamma waves (**Fig. 7c,d**). Our LFP analysis focused on computing Power Spectra Densities (PSDs) at these frequencies, with ranges selected from human brain wave investigation.

**Figure 7:**
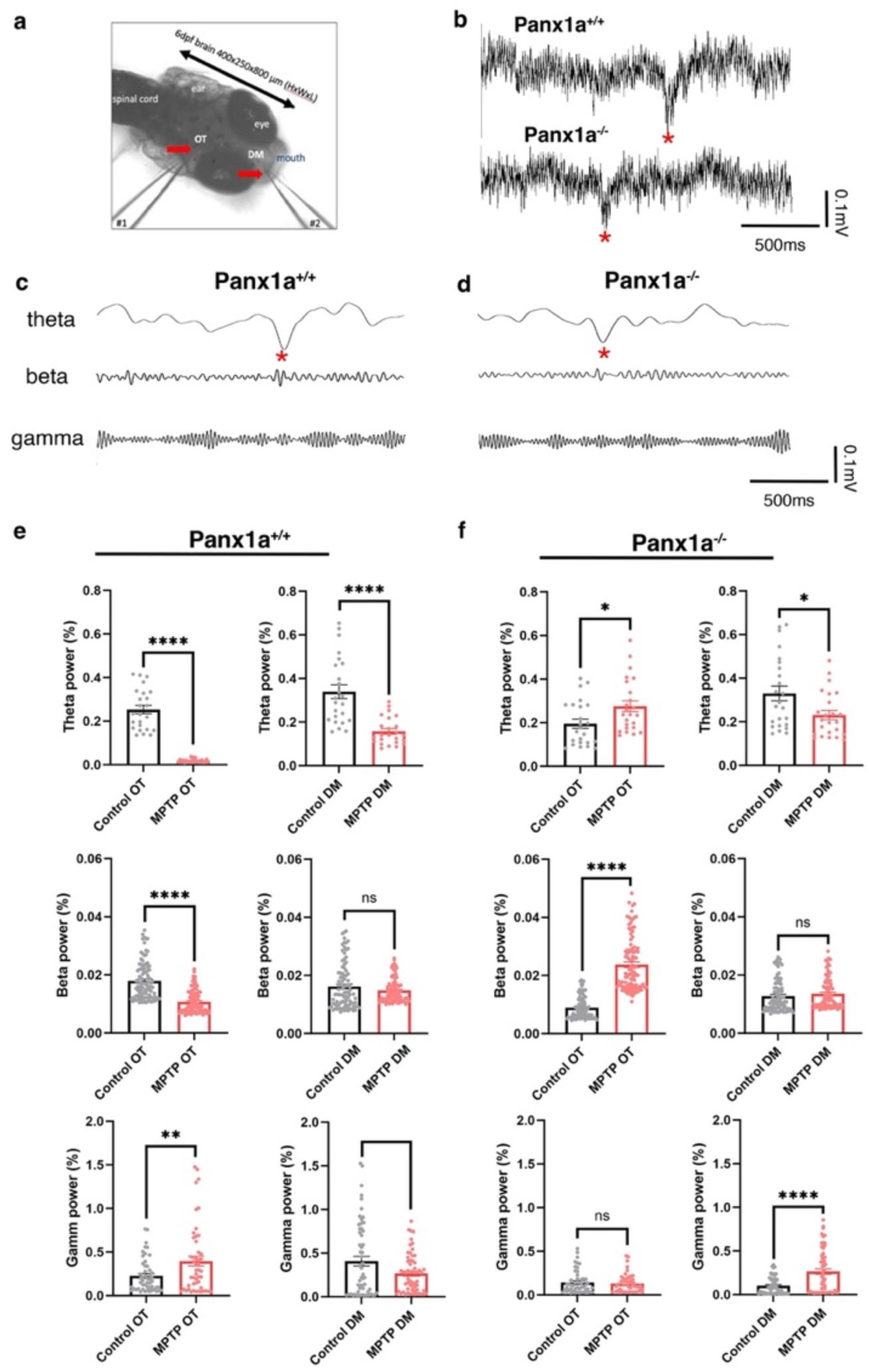
Loss-of-Panx1a function alters the power of theta, beta-and gamma-waves in the optic tectum. *a)* Representative image of a two-electrode setup with electrode #1 and electrode#2 placed into the optic tectum (OT) and dorsomedial forebrain (DM), respectively. b) Raw recording traces before isolating c,d) theta (3.5 – 7Hz), beta (12 -30Hz), and low gamma-wave (35 – 45Hz) frequency ranges. The red star indicates a reference point in the traces. e,f) Comparisons of the theta, beta, and gamma-power of Panx1a^+/+^ and Panx1a^-/-^ larvae without (control) and with 10µM MPTP treatment. Statistical approach: Outliers were removed using a ROUT test (Q = 1%). Significance was calculated using a Welch’s t-test. Number of larvae, n = 8 for Panx1a^+/+^ and Panx1a^-/-^. Mean ± SEM. Statistical significance was indicated as *: P-values * <0.05, ** <0.01, *** <0.005, **** <0.001, ns = not significant.

Significant inverse relationships were observed for theta, beta, and gamma power in the OTs of both genotypes (PSD % values: Theta -Panx1a^+/+^ 0.253 vs Panx1a^+/+MPTP^: 0.019, P-value < 0.0001, Panx1a^-/-^: 0.196 vs Panx1a^-/-MPTP^: 0.275, P-value = 0.0052; Beta - Panx1a^+/+^: 0.018 vs Panx1a^+/+MPTP^: 0.011, P-value < 0.0001, Panx1a^-/-^: 0.009 vs Panx1a^-/-MPTP^: 0.024, P-value < 0.0001; Gamma - Panx1a^+/+^: 0.227 vs Panx1a^+/+MPTP^: 0.397, P-value = 0.0012, Panx1a^-/-^: 0.145 vs Panx1a^-/-MPTP^: 0.131, P-value = 0.849). Similar trends were observed in the DM following MPTP treatment in both genotypes. PSDs of Panx1a^+/+^ and Panx1^-/-^ larvae either decreased in Panx1a^+/+^ (Theta: Panx1a^+/+^: 0.339 vs Panx1a^+/+MPTP^: 0.157, P-value < 0.0001, Panx1a^-/-^: 0.330 vs Panx1a^-/-MPTP^: 0.231, P-value = 0.188), remained unchanged (Beta: Panx1a^+/+^: 0.016 vs Panx1a^+/+MPTP^: 0.015, P-value = 0.660, Panx1a^-/-^: 0.013 vs Panx1a^-/-MPTP^: 0.014, P-value = 0.978), or decreased in Panx1a^+/+^ and increased in Panx1a^-/-^ (Gamma: Panx1a^+/+^: 0.407 vs Panx1a^+/+MPTP^: 0.267, P-value = 0.0465, Panx1a^-/-^: 0.101 vs Panx1a^-/-MPTP^: 0.264, P-value < 0.0001) (**Fig. 7e,f**).

We demonstrated that the LFPs in neuronal circuits in the OT and DM regions responded differently to MPTP+ when Panx1a was present. It was concluded that the changes occurring within selected spectral frequency bands connect functions of Panx1 to synaptic plasticity and mammalian CNS circuit pathophysiology.

Our findings demonstrate differential responses in LFPs within neuronal circuits of the OT and DM regions to MPTP when Panx1a is present. The observed changes in selected spectral frequency bands suggest a connection between Panx1 functions and synaptic plasticity, shedding light on mammalian CNS circuit pathophysiology.

### Loss-of-Panx1a reverses the coherence of local field potentials

A coherence analysis of LFPs in the frequency range from 0 to 60hz tested the impact of Panx1a ablation and MPTP treatment on the connectivity between the OT and DM, two pivotal regions in the ascending visual pathway (**Fig. 8a**). The zebrafish ascending visual pathway comprises polysynaptic connections, with retinal input to the OT and subsequent projections from the OT to the lateral preglomerular complex and the pallium, including the DM ^15,16^.

**Figure 8.**
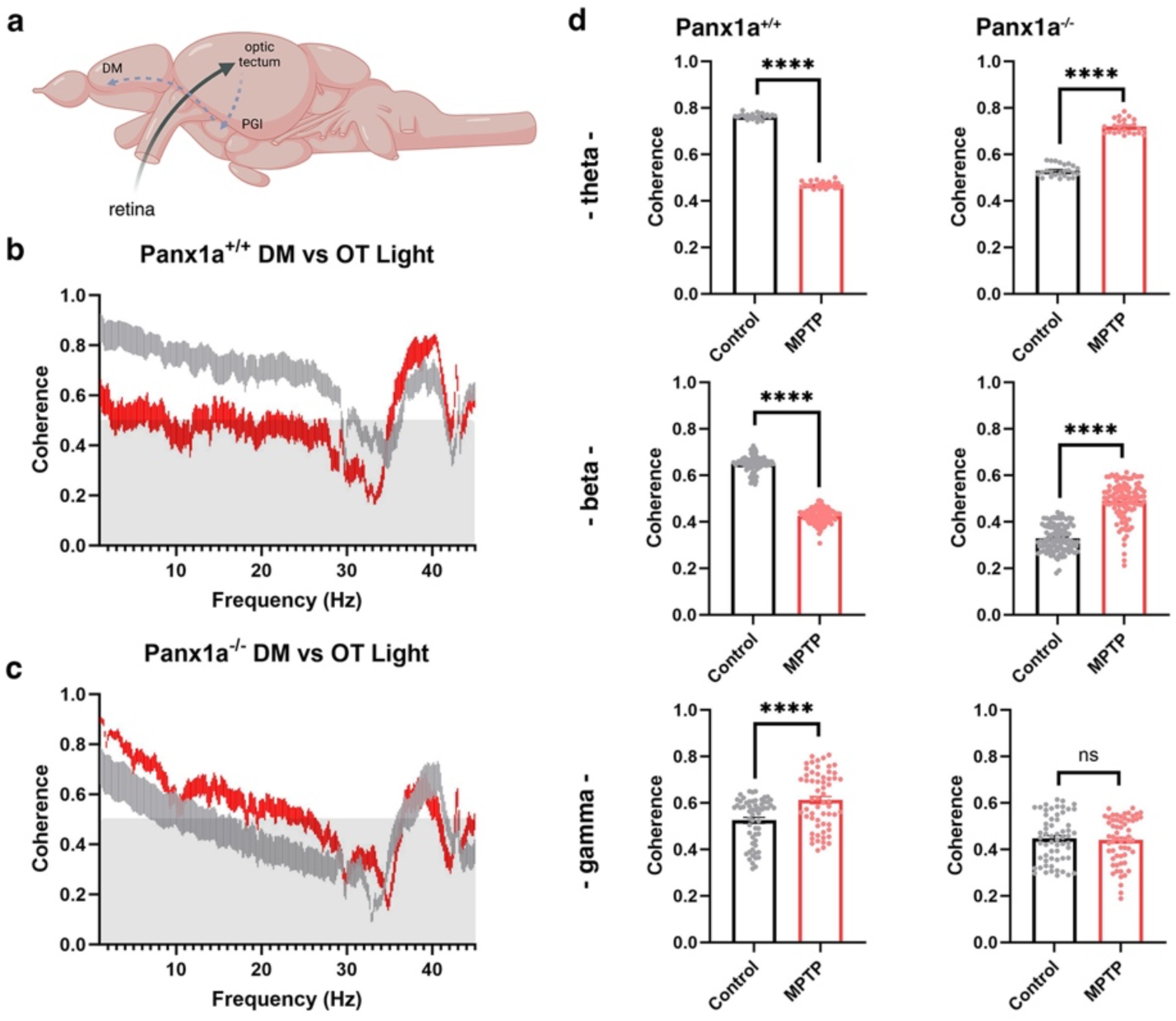
Loss-of-Panx1a function reverses the coherence of local field potentials. *a)* Outline of the ascending visual pathway. b,c) Shows the coherence of local field potentials measured in the optic tectum (OT) and the dorso-medial forebrain (DM) of 6dpf Panx1a^+/+^ and Panx1a^-/-^ larvae. The traces represent the 95% confidence intervals (CI) without (in grey) and after 4hrs of treatment with 50µM MPTP (in red). d) The theta (3.5 – 7Hz), beta (12 -30Hz) and gamma-waves (35 – 45Hz) coherence without (control, in grey) and after MPTP treatment (in red). Statistical approach: Outliers were removed using a ROUT test (Q = 1%). Significance was calculated using a Welch’s t-test. Number of larvae, n = 8 for Panx1a^+/+^ and Panx1a^-/-^. Mean ± SEM. Statistical significance was indicated as *: P-values * <0.05, ** <0.01, ***<0.005, **** < 0.001, ns= not significant.

MPTP treatment caused a reduction in coherence between OT and DM in Panx1a^+/+^ larvae across theta (Panx1a^+/+^: 0.764, Panx1a^+/+MPTP^: 0.470, P-value < 0.0001) and beta frequencies (Panx1a^+/+^: 0.650, Panx1a^+/+MPTP^: 0.425, P-value < 0.0001) (**Fig. 8b,c**). Panx1a^-/-^ larvae exhibited the converse effect, with augmented coherence in theta (Panx1a^-/-^: 0.523, Panx1a^-/-MPTP^: 0.469, P-value < 0.0001) and beta frequencies (Panx1a^-/-^: 0.330, Panx1a^-/-MPTP^: 0.490, P-value < 0.0001). Prior investigations have associated elevated beta frequency coherence with Parkinson’s disease pathology ^17^. The potential ramifications of the observed beta frequency coherence increase in Panx1a^-/-^ larvae warranted further exploration. Furthermore, Panx1a^+/+^ larvae demonstrated a notable increase in low gamma coherence (Panx1a^+/+^: 0.525, Panx1a^+/+MPTP^: 0.612, P-value 0.0001), a phenomenon previously linked to uncertainty ^18^.

Collectively, alterations in theta, beta, and low gamma coherence in Panx1a^+/+^ larvae suggested that MPTP treatment induced a functional modulation of neural network mechanisms, likely involving a combination of loss of cell homeostasis, synaptic plasticity, and cell death.

## Discussion

In this study acute MPTP-induced neurotoxicity served as a model to study Panx1a function in a period of zebrafish neurodevelopment that is considered to resemble late-term to birth in Mammalia. The principal results provide insights into how Panx1a affects the complex mechanisms of metabolic stress, including oxidative stress, mitochondrial dysfunction, and altered synaptic plasticity, thereby making inroads into the metabolic and synaptic aspects of neurodevelopmental and neurodegenerative diseases. Collectively we introduce a first-hand perspective on the role of Panx1a channels in the context of acute MPTP-induced changes of mitochondrial health in dopaminergic neurons; the evidence supports a role of Panx1a in regulating complex cell communication and signaling pathways required for the maintenance of neuronal homeostasis and network functions.

### Panx1a mediates neuronal homeostasis via AMPK-pathway activation and mTOR signaling

The upregulation of mitochondrial Complex I and III genes in Panx1a-deficient and MPTP treated larvae indicates significant changes in mitochondrial homeostasis. These changes are associated with pathways leading to hypoxia, reactive oxygen species (ROS) production, glucose and fatty acid metabolism, or the unfolded protein response, underscoring the necessity of Panx1a for maintaining metabolic and energetic balance *in vivo*.

Although the direct functions of Panx1 in mitochondrial health in the nervous system are not fully understood, Panx1a likely operates akin to a “rheostat.” In this concept, Panx1 channels open and close in response to varying brain states, with ATP and calcium concentration dynamics serving as messengers of these states. The known structural aspects of Panx1 proteins, such as functioning as a large pore ATP release channel or a narrow Cl channel ^19,20^, interacting with different lipid environments ^21^, or interacting proteins ^22^, could be contributing to a rheostat function.

The regulation of the AMP-activated protein kinase (AMPK) pathway showed the activation of a crucial cellular energy sensor to maintain energy homeostasis ^23^. AMPK is activated in response to an increase in the AMP/ATP ratio, which occurs during cellular stress or energy depletion ^24^. We identified both ATP depletion and the differential expression of subunits of the zebrafish AMPK in both genotypes after MPTP treatment suggesting simultaneous loss of Panx1-mediated ATP release and depletion of ATP after Complex I blocking can activate AMPK to enhance alternative energy production pathways.

Alternative catabolic pathways like the observed transcriptional activation of glycolysis and fatty acid metabolism help to restore the energy balance by inhibiting anabolic processes and while stimulating catabolic pathways ^25^ However, prolonged ATP depletion can trigger endoplasmic reticulum (ER) stress, initiating the unfolded protein response (UPR) ^26^. While the UPR seeks to restore protein homeostasis, severe or persistent ER and mitochondrial stress can activate apoptotic pathway ^27^. Interestingly, activation of Pannexin-1 and AMPK signaling pathway in the cerebral cortex of a mouse model of Sepsis-associated encephalopathy (SAE) was coupled with the activation of pyroptosis, a highly inflammatory form of lytic programmed cell death, and the incomplete activation of autophagy, which breaks down and recycles old, damaged, or abnormal proteins, especially during periods of stress or starvation ^28^. In contrast, the inhibition of pannexin-1 expression reduced the rates of mortality and the cortical pathological changes in the mice, further stimulating the AMPK signaling pathway, inhibiting pyroptosis, and activating autophagy ^28^. Here, we observed an up-regulation of autophagy and apoptosis related genes but no differential expression of GasderminD/E expression, a hallmark of pyroptosis, was not detected. This observation supports a theory that by targeting Panx1 in mice and fish inflammation caused by neuronal pyroptosis can be regulated through autophagy activation by the AMPK signaling pathway.

In Panx1a^-/-MPTP^ larvae, intensified cell death via autophagy seems to represent an exit strategy in response to disturbed neuronal homeostasis and mitochondrial health. Our data implicate specific cell death pathways, notably those involving the mammalian target of rapamycin complex 1 (mTORC1). In general, mTOR regulates various cellular functions such as protein synthesis, energy metabolism, cell size, lipid metabolism, autophagy, and lysosome biogenesis ^29^. The mTOR-controlled signaling pathways are crucial for neuronal development, synaptic plasticity, memory storage, and cognition ^30^. Dysregulation of mTOR signaling is a common feature of neurological and neuropsychiatric disorders. Mutations in synaptic genes and mitochondrial perturbation is increasingly recognized as a significant driver of synaptic dysfunction in autism spectrum disorder (ASD) ^31^. Individuals with ASD often show deficits in mitochondrial oxidative phosphorylation, linking mitochondrial dysfunction to the disorder’s development ^32^. Recent research has highlighted the significant roles of AMP-activated protein kinase (AMPK) and the mammalian target of rapamycin (mTOR) signaling pathways in the pathogenesis of ASD ^33^. Genetic mutations in regulators of mTOR, such as TSC1, TSC2, and PTEN, have been associated with ASD-like phenotypes, suggesting that mTOR dysregulation contributes to the disorder’s development. Moreover, the PI3K/Akt/mTOR pathway’s overactivation during brain development can lead to increased cell death, neuroinflammation, and oxidative stress, further exacerbating ASD symptoms ^33^. AMPK, a key energy sensor in cells, interacts with the mTOR pathway, and its overactivation has been shown to suppress mTOR signaling, leading to defects in dendritic morphology and increased susceptibility to anxiety, which is often comorbid with ASD ^34^. Overall, the AMPK and mTOR pathways are crucial in the neurodevelopmental processes underlying ASD. Interestingly, two studies recently linked Panx1 to ASD; ASD susceptibility genes were found in the murine Panx1 interactome, and loss of Panx1 changes social behaviors in an experimental model for early-life seizures ^22,35^.

### Panx1a modulates the molecular composition and bioelectrical function of synapses

Information about Panx1’s involvement in transsynaptic activities and synapse organization was found in the differential expression of genes encoding proteins with pre-and postsynaptic localization in metabolically challenged zebrafish. The expression changes include the downregulation of the presynaptic SNARE complex, postsynaptic excitatory ionotropic AMPA-, NMDA-, and kainite receptors subunits, inhibitory GABA-gated chloride ion channel, or anion and cation channels. Other molecular changes include the downregulation of biomarkers affecting neuronal homeostasis, synapse function, and axonal processes. The downregulation of genes encoding both pre-and postsynaptic proteins suggest trans-synaptic functions of Panx1’s function associated with synaptic homeostasis in both excitatory and inhibitory neurons.

In 2007, we have postulated that Panx1 channel localization in postsynaptic contacts of mouse cerebral cortex and the hippocampus represents a strategic position for synapse function modulation ^36^. Our group and others later demonstrated that loss of murine Panx1 channels causes enhanced and functional modifications of synapses encompassing increased hippocampal excitability, complex dendritic branching, enhanced spine maturation, and an increased proportion of multiple synaptic contacts in hippocampal neurons of Panx1-KO mice ^37–39^. Further, Panx1-mediated currents were potentiated by metabotropic receptors and bidirectionally modulated by burst-timing-dependent plasticity of NMDAR-mediated transmission ^40^. Moreover, under genetic conditions associated with neurodegenerative disorders (NDDs), suppression of Panx1 can restore synaptic plasticity. For example, in a transgenic Alzheimer’s Disease model, pharmacological blocking of Panx1 significantly reduced excitatory synaptic defects by normalizing long-term potentiation (LTP) and depression (LTD) and improving dendritic arborization and spine density in hippocampal neurons ^41^. Genetic ablation of Panx1 suggested that Panx1 negatively regulated cortical dendritic spines development ^42^. In the visual system, Panx1 channels contribute to the synaptic modulation of a horizontal cell to a photoreceptor feedback mechanism in the horizontal cell dendrites of the mouse and the zebrafish ^43–45^. Furthermore, loss of Panx1 caused altered learning and memory in mice such as diminished cognitive functions including altered spatial reference memory, or spatial reversal learning ^37,39,46,47^.

Despite accumulating evidence for the roles of Panx1 in synaptic plasticity, mechanistic details of the cellular processes and molecular pathways have been emerging slowly. Internalization of Panx1 by ATP and P2X7-dependent endocytosis and degradation ^48,49^ and suppression of presynaptic glutamate release via NMDA receptors and Panx1 have been reported ^50^. Panx1 coupling to NMDA receptor functions via Src ^51^, STIM1/2 ^52^ or actin, and Rho GTPases, appear to be relevant for synapse stability ^38^. Panx1 channel blockage with the mimetic peptide ^10^Panx1 increases the synaptic level of endocannabinoids (eCB) and the activation of cannabinoid receptors type 1 (CB1Rs), decreasing hippocampal GABAergic efficacy and shifting excitation/inhibition (E/I) balance toward excitation and facilitating the induction of long-term potentiation ^53^. Further, we have reported a pathway that connects NMDA receptor signaling to the opening of electrical synapse protein Cx36 via Panx1 and CaMK2a, suggesting the active participation of Panx1 in modulating both chemical and electrical synapse function by using a shared molecular machinery ^54^.

Jointly, the new discoveries support Panx1’s crucial role in cell communication and signaling processes in neurons, emphasizing an association with synaptic stability, memory formation, and intercellular signaling. Here, we demonstrated that Panx1a ablation influenced local field potential power of theta, beta, and gamma waves in the optic tectum and dorsomedial forebrain. This result implies a correlation between metabolic changes and synapse functions providing insights into how Panx1a deficiency alters brain wave activity and neuronal circuit responses. The modulation of coherence is evidence that the loss of Panx1a leads to changes in the relationship of the different brain regions across various frequency bands. Together, the findings of the LFP analysis establish a link between Panx1a functions and synaptic plasticity within neuronal circuits, supporting a role of Panx1a in maintaining synaptic stability and modulating brain activity ultimately correlating changes of brain function with the differential expression of pre-and postsynaptic genes.

The results fill a gap in understanding how Panx1a affects neural network connectivity and information processing, offering insight into Panx1’s role in disease pathology. Indeed, local field potentials (LFPs) can be altered in neurodevelopmental disorders. In autism spectrum disorder (ASD), mice prenatally exposed to valproic acid (VPA) exhibited significant changes in LFPs across several brain regions ^55^. These mice showed increased low gamma activity in the hippocampus during awake immobility and altered theta oscillations correlated with locomotor speed, which were not observed in control mice. Additionally, increased delta and beta activities were found in the olfactory bulb during different states of activity, along with altered coherence between the hippocampus and both the olfactory bulb and medial prefrontal cortex, suggesting disrupted neural signaling in learning-related brain areas ^55^. Notably, LFPs reflect the activity of spatially localized populations of neurons, and alterations in LFP activity can be indicative of cognitive processes, such as attention, which may be disrupted in neurodevelopmental disorders.

Local field potentials (LFPs) are also altered in neurodegenerative disorders, reflecting changes in neural network activities and brain connectivity. In Alzheimer’s disease (AD), studies on APP/PS1 mice models have shown that the LFPs in the gamma band of the left and right secondary motor cortex (M2) are more synchronized compared to wild-type controls, indicating a decline in hemispheric asymmetry, which could be an early sign of AD ^56^. Similarly, in Parkinson’s disease (PD), alterations in LFPs have been observed in several brain regions ^57^. Beyond AD and PD, LFP alterations have also been noted in psychiatric conditions like major depressive disorder (MDD) and obsessive-compulsive disorder (OCD) ^58,59^. These studies have found that different frequency bands are associated with specific symptoms, with low-frequency activity closely related to OCD symptoms, while LFP findings in MDD are more complex and varied ^60^ ^61^.

## Conclusion

Overall, we show that Panx1a is crucial for neuronal viability by regulating energy metabolism, facilitating cellular communication, and maintaining synaptic homeostasis. Our research identifies the AMPK-mTORC1 pathway as a key regulator of processes activating cell death pathways that allow the orderly degradation and recycling of cellular components. We propose that the evidence strongly suggests that Panx1a acts as a safeguard for neuronal health by modulating ATP signaling, thereby preserving metabolic and energetic homeostasis. The critical role of Panx1a in regulating synaptic plasticity and neuronal homeostasis, is likely impacting neuropsychiatric disorders like autism spectrum disorder (ASD) or early in life disturbances leading to neurodegenerative disorders like Alzheimer’s or Parkinson’s disease.

## Acknowledgements

We thank two members of York University Zebrafish Vivarium, Janet Fleites-Medina, and Veronica Scavo for outstanding zebrafish husbandry. We also wish to thank the Center for Applied Genomics, SickKids, Toronto, ON, Canada for the RNA-seq service.

## Author contributions

Conceptualization, GSZ, GRZ; data analysis, GSZ, NS, GRZ; investigation, GSZ, NS, CZ; writing—original draft preparation, GSZ, GRZ; writing—review and editing, all authors; visualization, GSZ, NS, GRZ; supervision, SC, GRZ; project administration, GRZ; funding acquisition, GRZ.

## Consent for publication

All authors have read and agreed to the published version of the manuscript.

## Ethics approval

All animal work was performed at York University’s zebrafish vivarium and in an S2 biosafety laboratory following the Canadian Council for Animal Care guidelines after approval of the study protocol by the York University Animal Care Committee (GZ#2019-7-R2).

## Data Availability

The RNA-seq data are deposited at the NCBI -Gene Expression Omnibus (GEO) database repository (ID pending at time of submission). Additional information necessary for the reanalysis of the data reported in this manuscript is available from the corresponding author upon request.

## Materials & Correspondence

Correspondence and material requests should be addressed to Georg R. Zoidl.

*Gene IDs follow ZFIN Zebrafish Nomenclature standards:* https://zfin.atlassian.net/wiki/spaces/general/pages/1818394635/ZFIN+Zebrafish+Nomenclature+Conventions

## Abbreviations

AD: Alzheimer’s Disease
ALS: Amyotrophic lateral sclerosis
AMPA: α-amino-3-hydroxy-5-methyl-4-isoxazolepropionic acid
AO: Acridine-Orange
ATP: Adenosine triphosphate
CNS: Central nervous system
DM: Dorsomedial pallium (amygdala)
DGE: differential gene expression
dpf: days post fertilization
FDR: False discovery rate
FFT: fast Fourier transform
Fps: frames per second
GABA: γ-Aminobutyric acid
GEO: Gene expression omnibus
GO: Gene Ontology
GSEA: Gene set enrichment analysis
HD: Huntington’s Disease
KEGG: Kyoto Encyclopedia of Genes and Genomes
LFP: Local field potential
NMDA: N-Methyl-D-aspartic acid
MSigDB: Molecular Signatures Database
MPTP: 1-methyl-4-phenyl-1,2,3,6-tetrahydropyridine
NDD: neurodegenerative disorder
OT: Optic tectum
Padj: adjusted P-value
Panc: Pancuronium bromide
PD: Parkinson’s Disease
PCA: Principal component analysis
PSD: Power spectral density
PROBS: Phosphate-Buffered Saline
ROS: Reactive oxygen species
RNA-seq: RNA sequencing
SEM: standard error of the mean
SNARE: soluble N-ethylmaleimide-sensitive fusion protein attachment protein receptors
TL: Tubingen longfin
UPR: Unfolded protein response
ZFIN: Zebrafish Information Network

## References

1 Safarian, N., Whyte-Fagundes, P., Zoidl, C., Grigull, J. & Zoidl, G. Visuomotor deficiency in panx1a knockout zebrafish is linked to dopaminergic signaling. Sci Rep 10, 9538 (2020). 10.1038/s41598-020-66378-y

2 Safarian N., H.-T. S., Zoidl C., Zoidl G.R. Panx1b modulates the luminance response and direction of motion in the zebrafish. BioRxiv **BIORXIV-2021-453251v1-Zoidl** (2021).

3 Safarian, N., Houshangi-Tabrizi, S., Zoidl, C., & Zoidl, G.R. Panx1b Modulates the Luminance Response and Direction of Locomotion in the Zebrafish. Int. J. Mol. Sci. (2021). 10.3390/ijms222111750

4 MacPhail, R. C. et al. Locomotion in larval zebrafish: Influence of time of day, lighting and ethanol. Neurotoxicology 30, 52–58 (2009). 10.1016/j.neuro.2008.09.011

5 Safarian, N., Houshangi-Tabrizi, S., Zoidl, C. & Zoidl, G. R. Panx1b Modulates the Luminance Response and Direction of Locomotion in the Zebrafish. Int J Mol Sci 22 (2021). 10.3390/ijms222111750

6 Baraban, S. C. Forebrain electrophysiological recording in larval zebrafish. J Vis Exp (2013). 10.3791/50104

7 Whyte-Fagundes, P. et al. A Potential Compensatory Role of Panx3 in the VNO of a Panx1 Knock Out Mouse Model. Front Mol Neurosci 11, 135 (2018). 10.3389/fnmol.2018.00135

8 McCarthy, D. J., Chen, Y. & Smyth, G. K. Differential expression analysis of multifactor RNA-Seq experiments with respect to biological variation. Nucleic Acids Res 40, 4288–4297 (2012). 10.1093/nar/gks042

9 Robinson, M. D., McCarthy, D. J. & Smyth, G. K. edgeR: a Bioconductor package for differential expression analysis of digital gene expression data. Bioinformatics 26, 139–140 (2010). 10.1093/bioinformatics/btp616

10 Abad Tan, S., Zoidl, G. & Ghafar-Zadeh, E. A Multidisciplinary Approach Toward High Throughput Label-Free Cytotoxicity Monitoring of Superparamagnetic Iron Oxide Nanoparticles. Bioengineering (Basel) 6 (2019). 10.3390/bioengineering6020052

11 Seal, R. L. et al. Genenames.org: the HGNC resources in 2023. Nucleic Acids Res 51, D1003–D1009 (2023). 10.1093/nar/gkac888

12 Mudunuri, U., Che, A., Yi, M. & Stephens, R. M. bioDBnet: the biological database network. Bioinformatics 25, 555–556 (2009). 10.1093/bioinformatics/btn654

13 Ray, A., Zoidl, G., Weickert, S., Wahle, P. & Dermietzel, R. Site-specific and developmental expression of pannexin1 in the mouse nervous system. Eur J Neurosci 21, 3277–3290 (2005). 10.1111/j.1460-9568.2005.04139.x

14 Subramanian, A. et al. Gene set enrichment analysis: a knowledge-based approach for interpreting genome-wide expression profiles. Proc Natl Acad Sci U S A 102, 15545–15550 (2005). 10.1073/pnas.0506580102

15 Bloch, S. et al. Non-thalamic origin of zebrafish sensory nuclei implies convergent evolution of visual pathways in amniotes and teleosts. Elife 9 (2020). 10.7554/eLife.54945

16 Mueller, T. What is the Thalamus in Zebrafish? Front Neurosci 6, 64 (2012). 10.3389/fnins.2012.00064

17 Asadi, A., Madadi Asl, M., Vahabie, A. H. & Valizadeh, A. The Origin of Abnormal Beta Oscillations in the Parkinsonian Corticobasal Ganglia Circuits. Parkinsons Dis 2022, 7524066 (2022). 10.1155/2022/7524066

18 Nurislamova, Y. M., Novikov, N. A., Zhozhikashvili, N. A. & Chernyshev, B. V. Enhanced Theta-Band Coherence Between Midfrontal and Posterior Parietal Areas Reflects Post-feedback Adjustments in the State of Outcome Uncertainty. Front Integr Neurosci 13, 14 (2019). 10.3389/fnint.2019.00014

19 Dahl, G. The Pannexin1 membrane channel: distinct conformations and functions. FEBS Lett 592, 3201–3209 (2018). 10.1002/1873-3468.13115

20 Mim, C., Perkins, G. & Dahl, G. Structure versus function: Are new conformations of pannexin 1 yet to be resolved? J Gen Physiol 153 (2021). 10.1085/jgp.202012754

21 Anderson, C. L. & Thompson, R. J. Intrapore lipids hydrophobically gate pannexin-1 channels. Sci Signal 15, eabn2081 (2022). 10.1126/scisignal.abn2081

22 Frederiksen, S. D., Wicki-Stordeur, L. E. & Swayne, L. A. Overlap in synaptic neurological condition susceptibility pathways and the neural pannexin 1 interactome revealed by bioinformatics analyses. Channels (Austin) 17, 2253102 (2023). 10.1080/19336950.2023.2253102

23 Muraleedharan, R. & Dasgupta, B. AMPK in the brain: its roles in glucose and neural metabolism. FEBS J 289, 2247–2262 (2022). 10.1111/febs.16151

24 Steinberg, G. R. & Hardie, D. G. New insights into activation and function of the AMPK. Nat Rev Mol Cell Biol 24, 255–272 (2023). 10.1038/s41580-022-00547-x

25 Athari, S. Z., Farajdokht, F., Keyhanmanesh, R. & Mohaddes, G. AMPK Signaling Pathway as a Potential Therapeutic Target for Parkinson’s Disease. Adv Pharm Bull 14, 120–131 (2024). 10.34172/apb.2024.013

26 Volgyi, K., Juhasz, G., Kovacs, Z. & Penke, B. Dysfunction of Endoplasmic Reticulum (ER) and Mitochondria (MT) in Alzheimer’s Disease: The Role of the ER-MT Cross-Talk. Curr Alzheimer Res 12, 655–672 (2015). 10.2174/1567205012666150710095035

27 Ekundayo, B. E. et al. Oxidative Stress, Endoplasmic Reticulum Stress and Apoptosis in the Pathology of Alzheimer’s Disease. Cell Biochem Biophys 82, 457–477 (2024). 10.1007/s12013-024-01248-2

28 Lei, Y. et al. The pannexin-1 channel regulates pyroptosis through autophagy in a mouse model of sepsis-associated encephalopathy. Ann Transl Med 9, 1802 (2021). 10.21037/atm-21-6579

29 Panwar, V. et al. Multifaceted role of mTOR (mammalian target of rapamycin) signaling pathway in human health and disease. Signal Transduct Target Ther 8, 375 (2023). 10.1038/s41392-023-01608-z

30 Bockaert, J. & Marin, P. mTOR in Brain Physiology and Pathologies. Physiol Rev 95, 1157–1187 (2015). 10.1152/physrev.00038.2014

31 Rojas-Charry, L., Nardi, L., Methner, A. & Schmeisser, M. J. Abnormalities of synaptic mitochondria in autism spectrum disorder and related neurodevelopmental disorders. J Mol Med (Berl) 99, 161–178 (2021). 10.1007/s00109-020-02018-2

32 Khaliulin, I., Hamoudi, W. & Amal, H. The multifaceted role of mitochondria in autism spectrum disorder. Mol Psychiatry (2024). 10.1038/s41380-024-02725-z

33 Thomas, S. D., Jha, N. K., Ojha, S. & Sadek, B. mTOR Signaling Disruption and Its Association with the Development of Autism Spectrum Disorder. Molecules 28 (2023). 10.3390/molecules28041889

34 Ma, J. et al. Dysregulation of AMPK-mTOR signaling leads to comorbid anxiety in Dip2a KO mice. Cereb Cortex 33, 4977–4989 (2023). 10.1093/cercor/bhac393

35 Obot, P. et al. Pannexin1 Mediates Early-Life Seizure-Induced Social Behavior Deficits. ASN Neuro 16, 2371164 (2024). 10.1080/17590914.2024.2371164

36 Zoidl, G. et al. Localization of the pannexin1 protein at postsynaptic sites in the cerebral cortex and hippocampus. Neuroscience 146, 9–16 (2007). 10.1016/j.neuroscience.2007.01.061

37 Ardiles, A. O. et al. Pannexin 1 regulates bidirectional hippocampal synaptic plasticity in adult mice. Front Cell Neurosci 8, 326 (2014). 10.3389/fncel.2014.00326

38 Flores-Munoz, C. et al. The Long-Term Pannexin 1 Ablation Produces Structural and Functional Modifications in Hippocampal Neurons. Cells 11 (2022). 10.3390/cells11223646

39 Prochnow, N. et al. Pannexin1 stabilizes synaptic plasticity and is needed for learning. PLoS One 7, e51767 (2012). 10.1371/journal.pone.0051767

40 Rangel-Sandoval, C., Soula, M., Li, W. P., Castillo, P. E. & Hunt, D. L. NMDAR-mediated activation of pannexin1 channels contributes to the detonator properties of hippocampal mossy fiber synapses. iScience 27, 109681 (2024). 10.1016/j.isci.2024.109681

41 Flores-Munoz, C. et al. Acute Pannexin 1 Blockade Mitigates Early Synaptic Plasticity Defects in a Mouse Model of Alzheimer’s Disease. Front Cell Neurosci 14, 46 (2020). 10.3389/fncel.2020.00046

42 Sanchez-Arias, J. C. et al. Pannexin 1 Regulates Network Ensembles and Dendritic Spine Development in Cortical Neurons. eNeuro 6 (2019). 10.1523/ENEURO.0503-18.2019

43 Vroman, R. et al. Extracellular ATP hydrolysis inhibits synaptic transmission by increasing ph buffering in the synaptic cleft. PLoS Biol 12, e1001864 (2014). 10.1371/journal.pbio.1001864

44 Kranz, K. et al. Expression of Pannexin1 in the outer plexiform layer of the mouse retina and physiological impact of its knockout. J Comp Neurol 521, 1119–1135 (2013). 10.1002/cne.23223

45 Cenedese, V. et al. Pannexin 1 Is Critically Involved in Feedback from Horizontal Cells to Cones. Front Mol Neurosci 10, 403 (2017). 10.3389/fnmol.2017.00403

46 Gajardo, I. et al. Lack of Pannexin 1 Alters Synaptic GluN2 Subunit Composition and Spatial Reversal Learning in Mice. Front Mol Neurosci 11, 114 (2018). 10.3389/fnmol.2018.00114

47 Obot, P. et al. Astrocyte and Neuronal Panx1 Support Long-Term Reference Memory in Mice. ASN Neuro 15, 17590914231184712 (2023). 10.1177/17590914231184712

48 Boyce, A. K., Kim, M. S., Wicki-Stordeur, L. E. & Swayne, L. A. ATP stimulates pannexin 1 internalization to endosomal compartments. Biochem J 470, 319–330 (2015). 10.1042/BJ20141551

49 Boyce, A. K. J. & Swayne, L. A. P2X7 receptor cross-talk regulates ATP-induced pannexin 1 internalization. Biochem J 474, 2133–2144 (2017). 10.1042/BCJ20170257

50 Bialecki, J. et al. Suppression of Presynaptic Glutamate Release by Postsynaptic Metabotropic NMDA Receptor Signalling to Pannexin-1. J Neurosci 40, 729–742 (2020). 10.1523/JNEUROSCI.0257-19.2019

51 Weilinger, N. L. et al. Metabotropic NMDA receptor signaling couples Src family kinases to pannexin-1 during excitotoxicity. Nat Neurosci 19, 432–442 (2016). 10.1038/nn.4236

52 Patil, C. S. et al. ER-resident STIM1/2 couples Ca(2+) entry by NMDA receptors to pannexin-1 activation. Proc Natl Acad Sci U S A 119, e2112870119 (2022). 10.1073/pnas.2112870119

53 Garcia-Rojas, F. et al. Pannexin-1 Modulates Inhibitory Transmission and Hippocampal Synaptic Plasticity. Biomolecules 13 (2023). 10.3390/biom13060887

54 Siu, R. C. F., Kotova, A., Timonina, K., Zoidl, C. & Zoidl, G. R. Convergent NMDA receptor-Pannexin1 signaling pathways regulate the interaction of CaMKII with Connexin-36. Commun Biol 4, 702 (2021). 10.1038/s42003-021-02230-x

55 Cheaha, D., Bumrungsri, S., Chatpun, S. & Kumarnsit, E. Characterization of in utero valproic acid mouse model of autism by local field potential in the hippocampus and the olfactory bulb. Neurosci Res 98, 28–34 (2015). 10.1016/j.neures.2015.04.006

56 Chen, Y., Li, M., Zheng, Y. & Yang, L. Evaluation of Hemisphere Lateralization with Bilateral Local Field Potential Recording in Secondary Motor Cortex of Mice. J Vis Exp (2019). 10.3791/59310

57 Wang, J. et al. Network-wide oscillations in the parkinsonian state: alterations in neuronal activities occur in the premotor cortex in parkinsonian nonhuman primates. J Neurophysiol 117, 2242–2249 (2017). 10.1152/jn.00011.2017

58 Trenado, C. et al. Combined Invasive Subcortical and Non-invasive Surface Neurophysiological Recordings for the Assessment of Cognitive and Emotional Functions in Humans. J Vis Exp (2016). 10.3791/53466

59 Zhang, W., Xiong, B., Wu, Y., Xiao, L. & Wang, W. Local field potentials in major depressive and obsessive-compulsive disorder: a frequency-based review. Front Psychiatry 14, 1080260 (2023). 10.3389/fpsyt.2023.1080260

60 Mitiureva, D. et al. Comparative analysis of resting-state EEG functional connectivity in depression and obsessive-compulsive disorder. Psychiatry Res Neuroimaging 342, 111828 (2024). 10.1016/j.pscychresns.2024.111828

61 Gimenez, M. et al. Brain alterations in low-frequency fluctuations across multiple bands in obsessive compulsive disorder. Brain Imaging Behav 11, 1690–1706 (2017). 10.1007/s11682-016-9601-y

